# Spatial Flavonoid Accumulation in Soybean Pericycle Restricts *Phytophthora sojae* Invasion

**DOI:** 10.64898/2025.11.30.691358

**Authors:** Qiushi Wang, Zhimin Dong, Haoyu Wang, Zhirui Zhang, Chunyang Ma, Xinwei Tan, Qunqing Wang, Yu Geng, Jiuhai Zhao

## Abstract

Root immunity relies on coordinated defenses across distinct tissues, yet the underlying cellular mechanisms remain unclear. Here, we integrated single-cell transcriptomics, spatial metabolomics, and genetic manipulation to dissect soybean (*Glycine max*) interactions with *Phytophthora sojae*. In resistant Williams82 (Wm82) roots, *P. sojae* hyphae were excluded from the stele, exhibiting negative tropism toward the pericycle. scRNA-seq revealed that this resistance is mediated by infection-induced pericycle cells expressing high levels of flavonoid biosynthetic genes, including *GmCHS7* and *GmCHI4A*. CRISPR-Cas9 knockout of these genes compromised resistance, whereas overexpression enhanced immunity in susceptible cultivars. Metabolomic and spatial metabolic analyses identified naringenin, a flavonoid phytoalexin, as a key antimicrobial compound that specifically accumulates in pericycle-adjacent tissues of Wm82 during infection. Naringenin directly inhibited hyphal growth *in vitro* and enhanced resistance when applied exogenously. These findings uncover a spatially targeted immune strategy where pericycle-localized activation of flavonoid synthesis forms a chemical barrier to safeguard vascular integrity, highlighting pericycle-centered metabolic immunity as a target for engineering durable resistance.

## Introduction

The oomycete pathogen *P. sojae* is a major causal agent of *Phytophthora* stem and root rot (PSRR), inflicting annual global losses of 1-2 billion USD and posing one of the most severe constraints on soybean production (Tyler et al., 2006; Tyler, 2007; Bradley et al., 2021). It produces thick-walled oospores capable of surviving for years in the absence of a host (Schmitthenner, 1985). As a soil-borne pathogen, *P. sojae* initiates infection from the root zone. The oospores germinate into sporangia under flooded soil conditions, and release motile zoospores, which is the primary infective agent responsible for root invasion (Tyler, 2007; Yang et al., 2013). The infection process involves an intricate molecular dialogue at the root–soil interface, where zoospores are chemotactically attracted to soybean root, and the pathogen secretes a repertoire of effectors to manipulate host immunity and metabolism (Hale et al., 2023; Zhu et al., 2023). Root exudates, particularly the isoflavonoids daidzein and genistein attract zoospores (Morris et al., 1998). When the soybean immune system fails to restrict *P. sojae* invasion, the pathogen rapidly colonizes root tissues, resulting in seedling damping-off and root rot in mature plants. Previous studies on soybean immunity to *P. sojae* have predominantly focused on aboveground tissues, particularly through hypocotyl inoculation assays (Lin et al., 2022). To date, at least 40 *Rps* genes have been reported (Lin et al., 2022; McCoy et al., 2023). These *Rps* genes confer race-specific resistance, typically governed by a single dominant locus, and have been widely deployed in commercial cultivars for PSRR management. Most identified *Rps* genes are predicted to encode nucleotide-binding site leucine-rich repeat (NBS-LRR) proteins that recognize *P. sojae* effectors and activate effector-triggered immunity (ETI) (Hale et al., 2023). Unfortunately, only limited number of *Rps* genes, including *Rps1k* (Gao et al., 2005) and *Rps11* (Wang et al., 2021), have been cloned and functionally validated, and the signaling events downstream of Rps–effector recognition remain largely unresolved.

Corresponding to the *Rps*—mediated response, virulence profiles of *P. sojae* evolves rapidly, compromising the effectiveness of many commercially deployed *Rps* genes, with over 200 pathotypes identified worldwide (Sahoo et al., 2021; McCoy et al., 2023). Although, the Rps–effector molecular interactions are diverse, it was suggested that *Rps*-mediated immunity elicits common cellular outcomes. In susceptible hosts, *P. sojae* hyphae colonize root tissues progressively—from the epidermis to the cortex and ultimately into the vascular stele (Enkerli et al., 1997). By contrast, incompatible interactions mediated by *Rps* genes trigger a rapid and localized hypersensitive response within 4 hours post infection, confining the pathogen to outer root cell layers through programmed cell death and cell wall reinforcement (WARD et al., 1989; Enkerli et al., 1997). These observations indicate that effective *Rps*-mediated resistance relies on a common physical and physiological barrier in root that prevents *P. sojae* from establishing systemic infection. A mechanistic understanding of root immunity is therefore critical for uncovering the principles of soybean–oomycete interactions and for guiding the development of durable resistance strategies. Despite the central role of roots as the initial site of pathogen entry and immune activation, the molecular and cellular mechanisms underlying root immunity to *P. sojae* remain poorly defined.

To address this gap, we applied single-cell transcriptomics to dissect the tissue-specific immune responses of soybean roots during *P. sojae* infection in the context of *Rps1k*-mediated resistance. Our investigation was particularly motivated by a pivotal and poorly understood observation: in the resistant genotype Wm82, *P. sojae* hyphae proliferate in the outer tissues but are consistently excluded from the vascular stele, exhibiting a marked negative tropism toward the pericycle. This spatial restriction of the pathogen stands in stark contrast to the rapid vascular colonization seen in susceptible hosts and suggests that, beyond the early hypersensitive response in outer layers, a highly specialized defense mechanism operates at this critical cellular boundary. We therefore hypothesized that the pericycle and adjacent tissues are not passive bystanders but active contributors to immunity, deploying a hitherto uncharacterized barrier that prevents stele entry. Furthermore, we hypothesized that distinct root cell layers respond differentially to *P. sojae* and that these responses constitute a downstream signaling cascade of *Rps*-mediated immunity. Indeed, our analyses revealed pronounced heterogeneity in immune activation across root cell types, with specific tissues exhibiting heightened defense activity—designated as “immune hotspots.” By single-cell RNA sequencing and spatial metabolomic analysis we demonstrated that pericycle-localized flavonoid biosynthesis is essential for *Rps1k*-mediated resistance. Overexpression of key flavonoid biosynthetic genes restricted *P. sojae* penetration into the stele, whereas application of a newly identified flavonoid phytoalexin naringenin significantly restored resistance in susceptible cultivars and conferred broad-spectrum protection against multiple *P. sojae* races. Together, these findings establish a mechanistic link between tissue-specific flavonoid metabolism and *Rps*-mediated root immunity, providing new insights into the spatial orchestration of defense responses against *P. sojae*.

## Results

### 1. *P. sojae* exhibits distinct growth patterns in compatible versus incompatible interactions with soybean roots

Previous studies have shown that during incompatible interactions between *P. sojae* and soybean roots, immune responses in the outer tissue layers—characterized by cell wall apposition and hypersensitive cell death—effectively block *P. sojae* colonization of the stele (WARD et al., 1989; Enkerli et al., 1997). However, the detailed dynamics of disease progression in compatible versus incompatible interactions remain unclear.

Here, we examined the susceptible soybean variety Williams (Wm) and its resistant derivative Williams82 (Wm82) (Figure 1A–C), which carries an resistance gene *Rps1k* (Gao et al., 2005).

**Figure 1.**
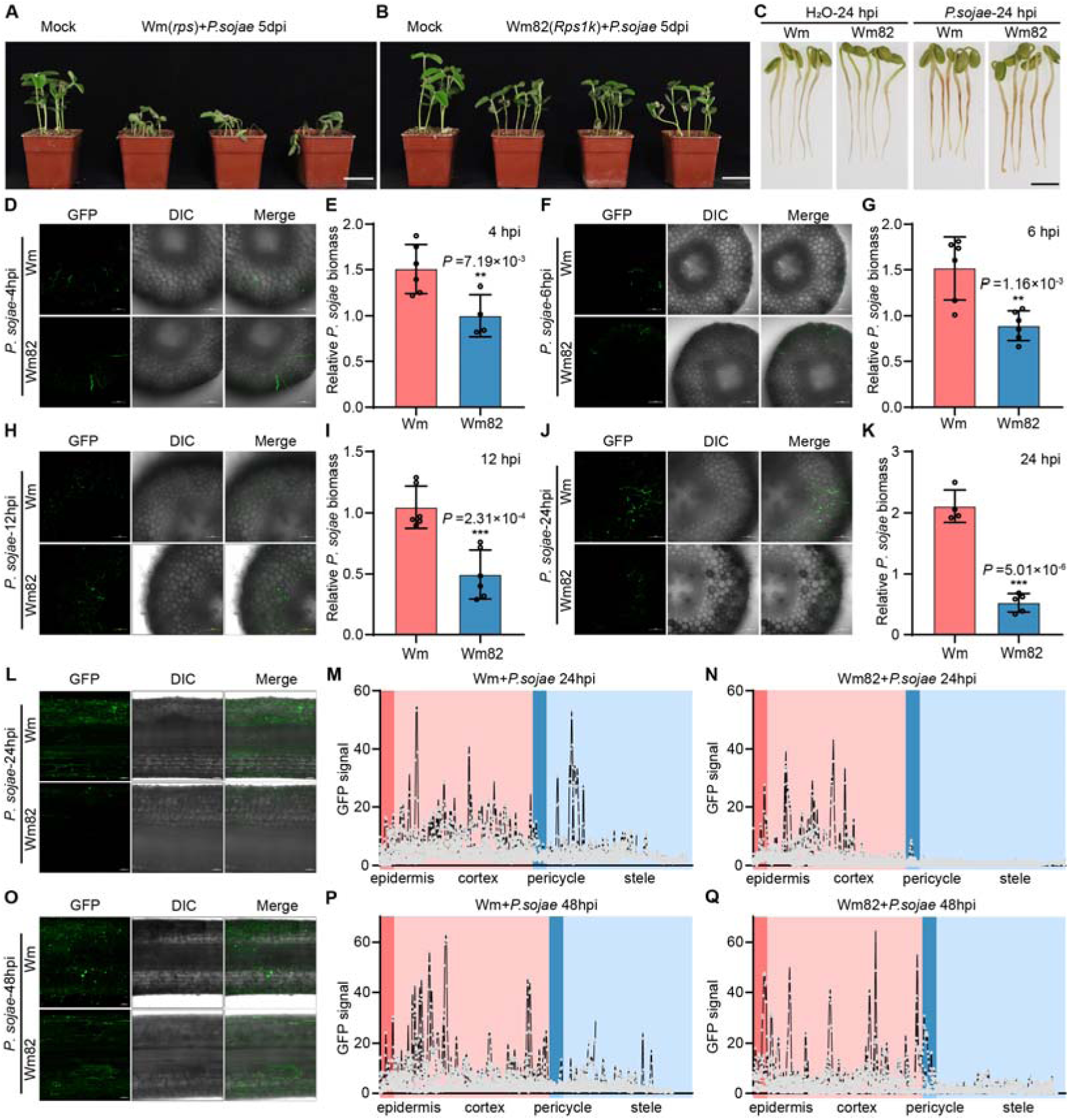
Infection outcomes and early colonization dynamics of *P. sojae* in susceptible Wm and resistant Wm82 soybean roots. (A, B) Disease symptoms of soil-grown Williams (Wm) and Williams 82 (Wm82) plants at 5 days post-inoculation (dpi) with *P. sojae*. Scale bar, 5 cm. (C) Etiolated Wm and Wm82 seedlings at 24 hours post-inoculation (hpi) with *P. sojae* zoospores or water (control). Scale bar, 1 cm. (D, F, H, J) Transverse root sections showing the progression of hyphal colonization in Wm and Wm82 at 4 h (D), 6 h (F), 12 h (H), and 24 h (J) post-inoculation. Scale bar, 100 μm. (E, G, I, K) Quantification of *P. sojae* biomass in Wm and Wm82 roots at 4 h (E), 6 h (G), 12 h (I), and 24 h (K) post-inoculation. Data represent means of three biological replicates, each with two technical replicates. **P ≤ 0.01; ***P ≤ 0.001. (L, O) Longitudinal root sections of Wm and Wm82 at 24 h (L) and 48 h (O) post-inoculation, illustrating contrasting colonization patterns between susceptible and resistant genotypes. (M, N, P, Q) GFP-tagged *P. sojae* colonization in Wm and Wm82 roots at 24 h (M, N) and 48 h (P, Q) post-inoculation. Data from three biological replicates.

Upon infection with *P. sojae* race 2, Wm displayed extensive root necrosis at 24 hours post-inoculation (hpi), whereas Wm82 showed minimal necrosis (Figure 1C). To compare pathogen growth over time, four-day-old etiolated seedlings of Wm and Wm82 were inoculated with GFP-labeled *P. sojae* zoospores, and disease progression was tracked using confocal imaging of free-hand sections from 4 to 48 hpi.

At 4 hpi, *P. sojae* hyphae were detected in the apoplast of cortical cells in both genotypes (Figure 1D). However, quantitative PCR (qPCR)-based biomass quantification revealed significantly higher pathogen loads in Wm compared to Wm82 (Figure 1E). This difference widened over time, as hyphal proliferation increased markedly in Wm but remained restricted in Wm82 (Figure 1F–K). By 24 hpi, hyphae had extensively colonized the stele in Wm but were absent from the stele in Wm82, where they instead grew along the pericycle (Figure 1L–N). Surprisingly, at 48 hpi, GFP signal intensity in Wm82’s outer layers was comparable to or higher than in Wm (Figure 1O), suggesting that the absence of stele colonization in Wm82 was not due to hyphal death in the peripheral tissues. Notably, hyphae in Wm82 exhibited negative tropism, avoiding stele penetration and aligning along the pericycle (Figure 1L).

These findings indicate a tissue-specific immune response in Wm82, potentially involving chemical or physical barriers around the stele that restrict *P. sojae* invasion. We propose that during incompatible interactions, distinct root tissues deploy specialized immune mechanisms to counter pathogen spread.

### 2. Single-cell transcriptomic atlas of soybean roots reveals early cell-type-specific responses to *P. sojae*

To investigate the tissue-specific aspects of the immune response to *P. sojae*, we performed high-throughput single-cell RNA sequencing (scRNA-seq) on samples from the Williams (Wm) and its resistant derivative Williams82 (Wm82).

Four-day-old etiolated seedlings of Wm and Wm82 were inoculated with *P. sojae* zoospores, with water-treated samples serving as controls (Figure S1). To investigate the early immune response and minimize the impact of infection on cell viability, root samples were collected and processed 4 hpi. Protoplasts were generated from the roots through enzymatic digestion using a cell wall-degrading enzyme cocktail (Ning et al., 2018). Single-cell transcriptome libraries were then prepared using the 10X Chromium platform. The gene expression matrices for each sample, processed with the Cellranger pipeline, underwent quality control in Seurat (Stuart et al., 2019). Low-quality cells and potential doublets were excluded, resulting in a final dataset of 8,189 high-quality cells from all four samples. These cells formed the foundation of our soybean root cell atlas.

Using an unbiased graph-based clustering approach in Seurat (Butler et al., 2018), we identified 15 distinct cell clusters. Cluster similarities, computed with Monocle2 (Trapnell et al., 2014) (Figure S2A), were used to guide cell type annotation. Primary annotations were performed with SCINA (Zhang et al., 2019) (Figure S2B, S2C) using marker genes curated from prior single-cell datasets (Table S1) (Shahan et al., 2022; Sun et al., 2023; Luo et al., 2024), pre-selected for cluster-specific expression in our cell atlas (Figure S3).

This annotation (Figure 2A) resolved all major root cell types, highlighting the robustness of our protoplast isolation and single-cell library construction. The expression of cluster-specific markers was provided in Figure 2B. Annotations were further validated and refined by in situ hybridization of cluster-specific genes. For example, *Glyma.11G095700* (*LOC100527864*), specifically expressed in C13 (initially annotated as stem cell niches), showed enriched expression in the stem cell niches and columella (Figure 2C). Similarly, *Glyma.15G040200* (*LOC100305705*), highly expressed in young endodermis and inner cortex, confirmed that C4 comprises meristematic ground tissue cells.

**Figure 2.**
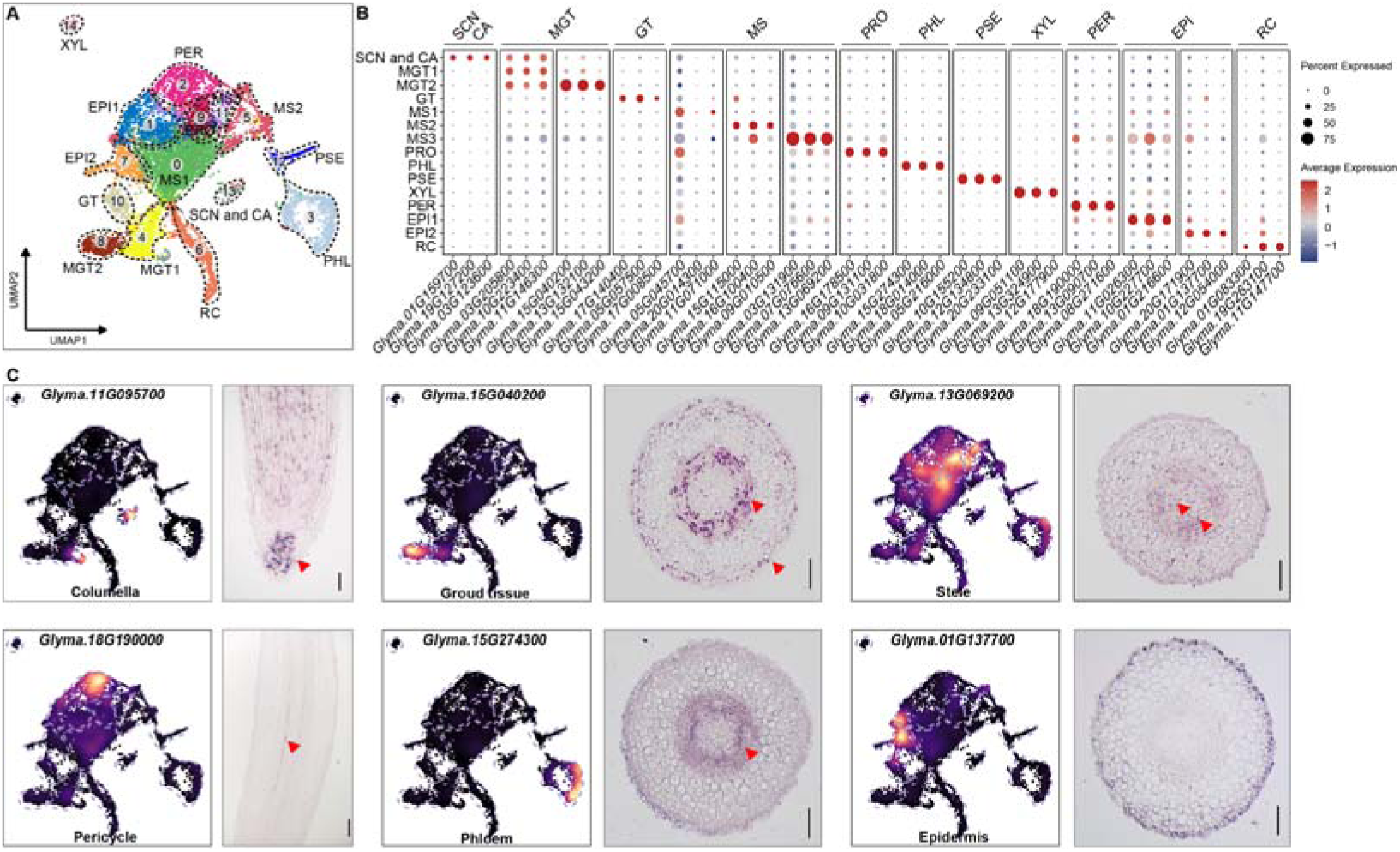
Construction and validation of a single-cell root atlas for Wm and Wm82 under *P. sojae* infection. (A) UMAP embedding of integrated single-cell transcriptomes from Wm and Wm82 roots under mock and *P. sojae* infection. Major cell types are annotated based on known tissue markers and transcriptional signatures. (SCN= stem cell niche; CA= columella; MGT= meristematic ground tissue; GT= ground tissue; MS= meristematic stele; PRO= procambium; PHL= phloem; PSE = phloem sieve elements; XLY= xylem; PER= pericycle; EPI= epidermis; RC= root cap) (B) Bubble plot showing representative cluster-specific marker genes across annotated cell types. Bubble size indicates the proportion of cells expressing each gene, and color intensity reflects scaled expression levels. (C) Validation of cell-type assignments using marker gene expression. Left: UMAP projection displaying expression patterns of selected marker genes. Right: corresponding in situ hybridization results in root sections confirming spatial localization of each marker. The concordance between in situ patterns and UMAP expression supports the accuracy of the cell atlas annotation. Scale bar, 100 μm.

### 3. Immune response landscapes differ markedly between compatible and incompatible interactions

To investigate the immune responses to *P. sojae* in both compatible and incompatible interactions, we conducted differential gene expression (DEG) analyses across distinct root cell types. To assess baseline differences between genotypes, we compared transcriptomes of Wm and Wm82 under control conditions. Only 26 genes were up-regulated and 40 down-regulated in Wm relative to Wm82, indicating minimal constitutive differences (Table S2).

We next compared infection-induced transcriptional changes between Wm and Wm82 across all cell clusters (Table S2; Fig. 3A,B). The resistant genotype Wm82 mounted a markedly stronger response, with 206 genes up-regulated and 166 down-regulated, compared with 128 and 177 in Wm, respectively. Notably, ∼70% of the Wm82-induced genes were enriched in the pericycle, highlighting this tissue as a major site of *Rps1k*-mediated defense activation. Only ∼13% of Wm82-upregulated genes were restricted to a single cluster, whereas 44% of infection-responsive genes in Wm showed cluster-specific induction, indicating a more spatially confined response in the susceptible genotype.

**Figure 3.**
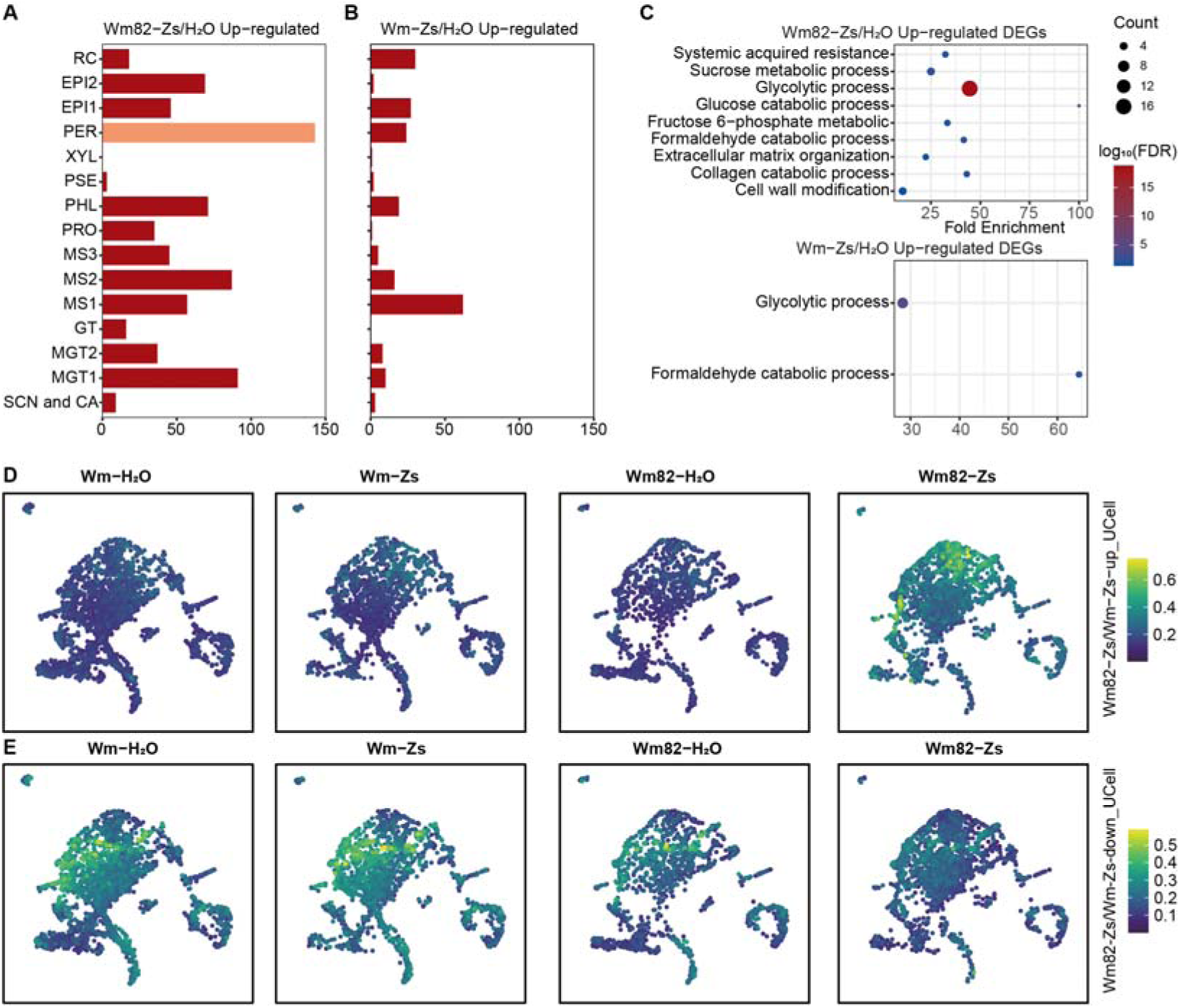
Genotype- and Cell-Type–Specific Transcriptional Reprogramming Highlights Pericycle-centered Defense Activation in Wm82. (A, B) Numbers of up-regulated genes across major root cell types in Wm and Wm82 after infection. The pericycle shows the largest increase in up-regulated genes in Wm82, whereas this pattern is not observed in Wm, supporting a pericycle-centered transcriptional response underlying resistance. (C) Gene Ontology (GO) enrichment analysis of differentially expressed genes (DEGs) between infected and control samples in Wm and Wm82. Wm82 exhibits stronger enrichment of pathways associated with central carbon catabolism, indicating enhanced metabolic activation and elevated energy demand during resistance. (D) UMAP projection of UCell scores for genes up-regulated in Wm82 relative to Wm following *P. sojae* infection, shown separately for each genotype and condition. The pericycle displays the strongest enrichment, indicating pronounced activation of infection-responsive programs in this tissue of the resistant genotype. (E) UMAP projection of UCell scores for genes down-regulated in Wm82 relative to Wm after infection. The epidermis displays the highest suppression signature, suggesting a transition of immune hotspot from epidermis to pericycle at the present of *Rps1k*.

GO enrichment analysis further revealed that Wm82 exhibited pronounced activation of central carbon catabolic pathways and systemic acquired resistance (SAR), suggesting elevated energy demand and metabolic reprogramming during resistance (Fig. 3C). These findings point to a broader and metabolically intensive defense program in Wm82.

We also identified 47 infection-induced genes shared between the two genotypes, but 39 of these were activated across a wider range of cell types in Wm82 than in Wm (Table S3; Fig. S4A). 11 shared DEGs belonged to glycolysis, further supporting enhanced energy mobilization in Wm82. Two expansin B-like genes (*Glyma.17G147400* and *Glyma.05G065700*) displayed restricted, cluster-specific activation in Wm yet were induced in 10 and 8 clusters, respectively, in Wm82. This expanded activation pattern is consistent with the known contribution of expansin B to resistance in other pathosystems (Brasileiro et al., 2021). UCell scoring of shared DEGs confirmed substantially higher expression levels in Wm82 (Fig. S4B), indicating that resistance involves both genotype-specific responses and stronger deployment of conserved defense genes.

To resolve the spatial organization of these transcriptional programs, we projected UCell enrichment scores of DEGs between Wm82 and Wm post infection onto the cell atlas. After infection, 192 genes were up-regulated and 109 down-regulated in Wm82 relative to Wm. Up-regulated genes localized predominantly to the pericycle, whereas down-regulated genes were concentrated in the epidermis (Fig. 3D,E). These transcriptionally dynamic tissues represent immune hotspots. The shift in immune hotspots—from epidermis-dominant in Wm to pericycle-dominant in Wm82—reflects a fundamental reorganization of defense signaling in the resistant genotype and likely underpins its ability to restrict *P. sojae* invasion.

### 4. Trajectory and coexpression analysis of Pericycle reveals connection between resistance and cell type-specific activation of flavonoid synthesis pathway

To further dissect the immune response to *P. sojae* within the pericycle, we performed pseudotemporal trajectory analysis using Monocle2 (Trapnell et al., 2014; Qiu et al., 2017). This analysis revealed a prominent branching event (branch point 4), which segregated pericycle cells into three transcriptionally distinct states (Figure S4). The pre-branch and cell type 2 groups contained cells from all four experimental conditions, indicating that a subset of pericycle cells maintain a stable transcriptional profile regardless of infection. In contrast, cell type 1 represented an infection-specific trajectory (Figure 4A). Cells from the susceptible genotype Wm were predominantly located at the early stages of this trajectory, whereas those from the resistant Wm82 genotype were enriched at later pseudotime points, suggesting a more extensive transcriptional reprogramming in response to infection (Figure 4B).

**Figure 4.**
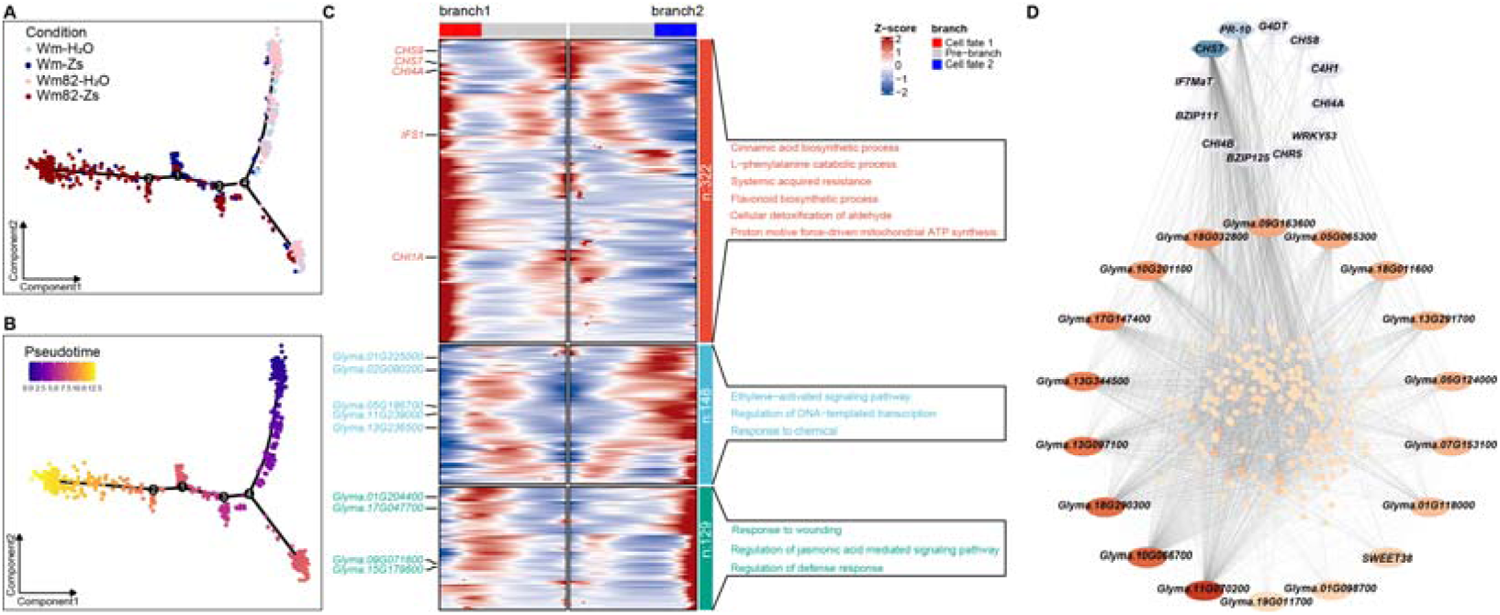
Pseudo-time analysis and WGCNA reveal distinct transcriptional dynamics in Wm82 pericycle cells during *P. sojae* infection and flavonoid biosynthesis activation. (A) Trajectory analysis of pericycle cells from all four conditions (*Wm-H_2_O*, *Wm-Zs*, *Wm82-H_2_O*, *Wm82-Zs*), showing a distinct branch predominantly comprising Wm82 post-infection cells. (B) Trajectory analysis of the pseudo-time across all samples. Cells from Wm82 post-infection occupy the highest pseudo-time values, reflecting more extensive transcriptional reprogramming during infection. (C) Heatmap showing significant genes associated with branch point 4, which divides the pericycle cells into three major branches, including the Wm82 post-infection branch and two branches encompassing all conditions. Gene clusters were identified by clustering branch-specific genes. Gene Ontology (GO) enrichment analysis for each gene cluster is shown, highlighting that flavonoid biosynthesis genes are enriched in the cluster highly expressed in Wm82 post-infection cells, suggesting an infection-induced activation of flavonoid metabolism. (D) Gene regulatory network (GRN) for the largest gene module identified by hdWGCNA analysis in pericycle cells. The network reveals coordinated expression of many flavonoid biosynthesis genes, underscoring their potential role in the resistance response.

To functionally characterize cell type 2, we performed gene clustering of branch-significant genes followed by Gene Ontology (GO) enrichment analysis (Yu et al., 2012). As shown in Figure 4C, three gene clusters were identified (Table S4), with cluster 3 genes—predominantly expressed in cell type 2—significantly enriched for biological processes related to systemic acquired resistance (SAR) and energy metabolism (Figure 4C, Table S5). These results support the interpretation that cell type 2 represents an immunologically active subpopulation, with Wm82-derived cells exhibiting a stronger immune activation signature than those from Wm.

To uncover potential regulatory mechanisms underlying this enhanced immune response, we conducted a co-expression network analysis using hdWGCNA (Morabito et al., 2023) for pericycle cluster. The largest gene module (Module Turquoise) showed strong correlation with Wm82 under infection (Figure S5). We extracted the genes from Module Turquoise and generated gene regulatory network with them. The resulting network (Figure 4D) identified key hub genes involved in flavonoid biosynthesis, including *CHS7*, *CHI4A* and *C4H1*. These flavonoid synthesis genes encode Flavonoids, as specialized metabolites, are well-known for their roles in plant defense against biotic stress (Treutter, 2005; Falcone Ferreyra et al., 2012; Mierziak et al., 2014). This finding raises the possibility that the activation of flavonoid biosynthesis contributes to the enhanced resistance phenotype observed in Wm82 during *P. sojae* infection.

### 5. Genetic manipulation of flavonoid synthesis pathway alters resistance to *P. sojae*

To further substantiate the link between the flavonoid synthesis pathway and resistance to *P. sojae*, we focused on the key infection-responsive flavonoid synthesis genes: *GmCHS7* and *GmCHI4A* (Figure S5), as these genes showed elevated expression in response to *P. sojae* specifically in Wm82. Our primary objective was to investigate whether genetic manipulation of these genes could influence the plant’s innate defense mechanisms against *P. sojae* infection. By utilizing both loss-of-function and gain-of-function approaches, we aimed to establish a causal relationship between the expression of these genes and resistance to this devastating pathogen.

To achieve this, we conducted CRISPR-Cas9 knockout experiments targeting each of the candidate genes using stable transformation system in the Wm82 genetic background. As illustrated in Figures 5A to 5C, and S6, disruption of any of these genes resulted in a significant increase in susceptibility to *P. sojae*. This heightened susceptibility was evidenced by more extensive pathogen colonization in the knockout roots compared to the control roots. Biomass assays revealed a notable increase in hyphal growth within the CRISPR knockout roots relative to the empty vector controls, suggesting a breakdown in the root’s capacity to mount an effective defense response. This finding strongly supports the hypothesis that these flavonoid synthesis genes are critical players in the plant’s immune defense. The elevated fungal biomass observed in knockout roots highlights the indispensable role of these genes in maintaining resistance.

**Figure 5.**
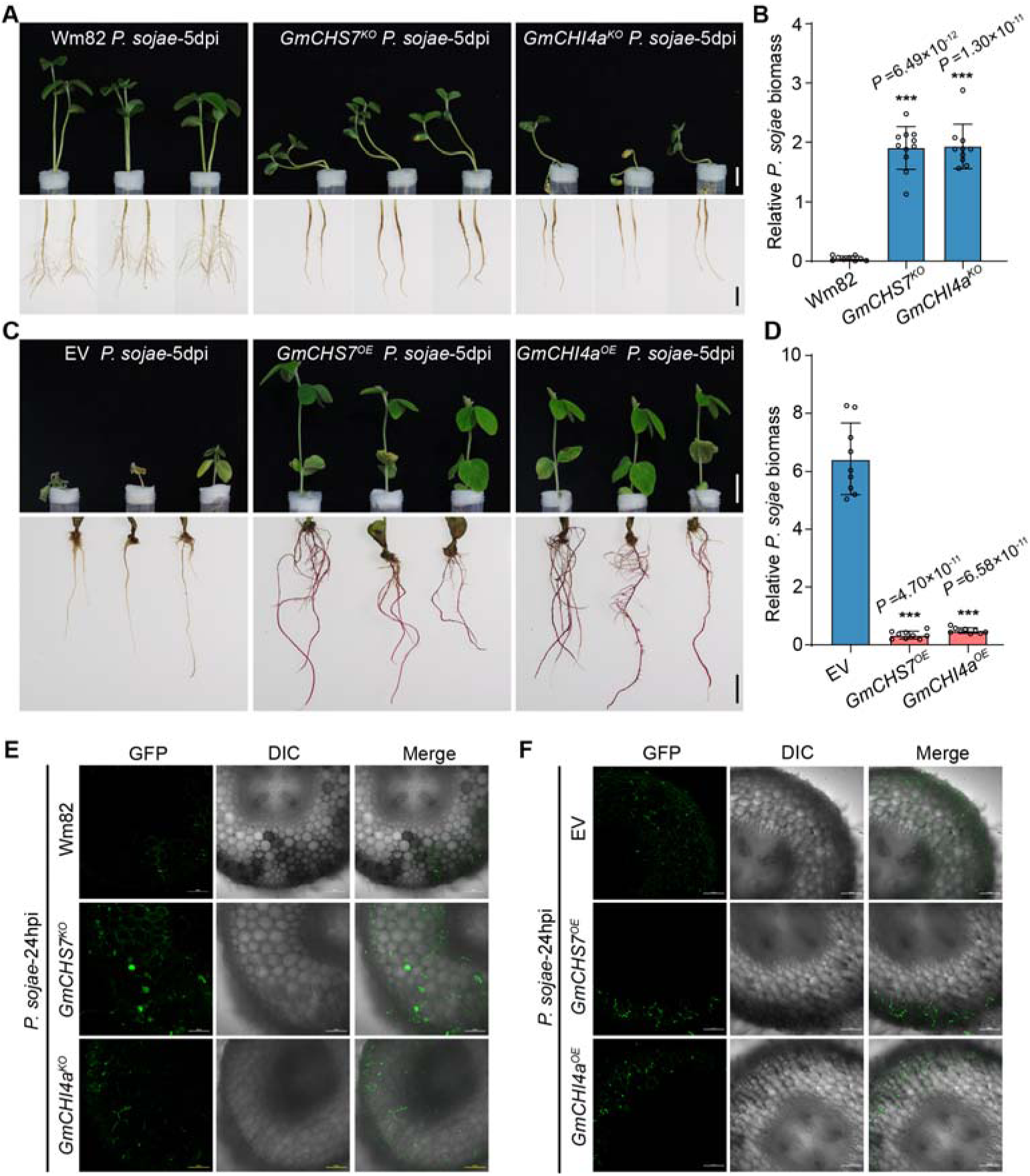
*GmCHS7* and *GmCHI4a* regulate flavonoid-dependent resistance to *P. sojae* invasion of the soybean root stele. (A) Root phenotypes of *GmCHS7* and *GmCHI4a* knockout (KO) transgenic plants at 5 days post-inoculation (dpi) with *P. sojae* zoospores. Loss of either gene compromises resistance and results in enhanced disease symptoms. Scale bar, 2 cm. (B) Relative *P. sojae* biomass in *GmCHS7^KO^* and *GmCHI4a^KO^* roots 4 hours post-inoculation (hpi). Both knockout lines show significantly increased pathogen accumulation compared with Wm82. (C) Root phenotypes of *GmCHS7* and *GmCHI4a* overexpression (OE) plants at 5 dpi with *P. sojae*. Overexpression enhances resistance and reduces disease severity. Scale bar, 2 cm. (D) Relative *P. sojae* biomass in *GmCHS7^OE^*and *GmCHI4a^OE^* roots 4 hpi. Overexpression markedly limits early pathogen growth compared with empty-vector controls. (E, F) GFP-labeled *P. sojae* infection in cross-sections of KO lines (E) and OE lines (F) at 24 hpi. Hyphae extensively colonize the stele in knockout plants but are strongly restricted in overexpression lines, demonstrating a key role for *GmCHS7* and *GmCHI4a* in preventing stele invasion. Scale bar, 100 μm.

Building on this insight, we sought to determine whether overexpression of these same flavonoid synthesis genes could confer enhanced resistance in a susceptible genetic background. To this end, we introduced the candidate genes into the susceptible Wm cultivar using a hairy root transformation system (Kereszt et al., 2007). As illustrated in Figures 5D to 5F, overexpression of *GmCHS7* and *GmCHI4A* significantly increased resistance against *P. sojae*, as reflected by lower pathogen biomass. These results are consistent with previous studies (Subramanian et al., 2005; Treutter, 2005; Falcone Ferreyra et al., 2012; Mierziak et al., 2014), that have highlighted the importance of the flavonoid pathway in plant immunity. Moreover, the data indicate that upregulating the expression of these flavonoid synthesis genes is sufficient to trigger enhanced defense responses, underscoring the feasibility of genetic engineering as a strategy for developing *P. sojae*-resistant soybean cultivars.

### 6. The Flavonoid synthesis pathway directly contribute to resistance against *P. sojae*

To investigate the role of flavonoid pathway products in conferring resistance to *P. sojae* in Wm82, we conducted a comprehensive metabolomics analysis (Sumner et al., 2015). This study included Wm, Wm82, and CRISPR knockout and overexpression lines targeting candidate flavonoid biosynthesis genes. The analysis was performed under both control conditions and at 24 hours post-infection.

From a targeted library of 3,500 flavonoid compounds, 604 were detected in our samples (Table S6). To investigate the relationship between these flavonoids and resistance to *P. sojae*, we conducted Pearson correlation analysis between metabolite levels and disease susceptibility across the Wm, Wm82, and CRISPR knockout lines of the candidate genes *GmC4H1*, *GmCHS7*, and *GmCHI4A* (Figure 6A). Overexpression lines were excluded from this analysis, as the strong ubiquitous expression of *GmC4H1*, *GmCHS7*, and *GmCHI4A* not only promoted the production of infection-related flavonoids but also generated numerous downstream products of the encoded enzymes (Table S6), thereby introducing substantial noise into the assay. We identified 192 metabolites that exhibited significant correlation (p < 0.05) with *P. sojae* biomass. Of these, 130 displayed positive correlations, while 62 showed negative correlations with pathogen biomass (Table S7).

**Figure 6.**
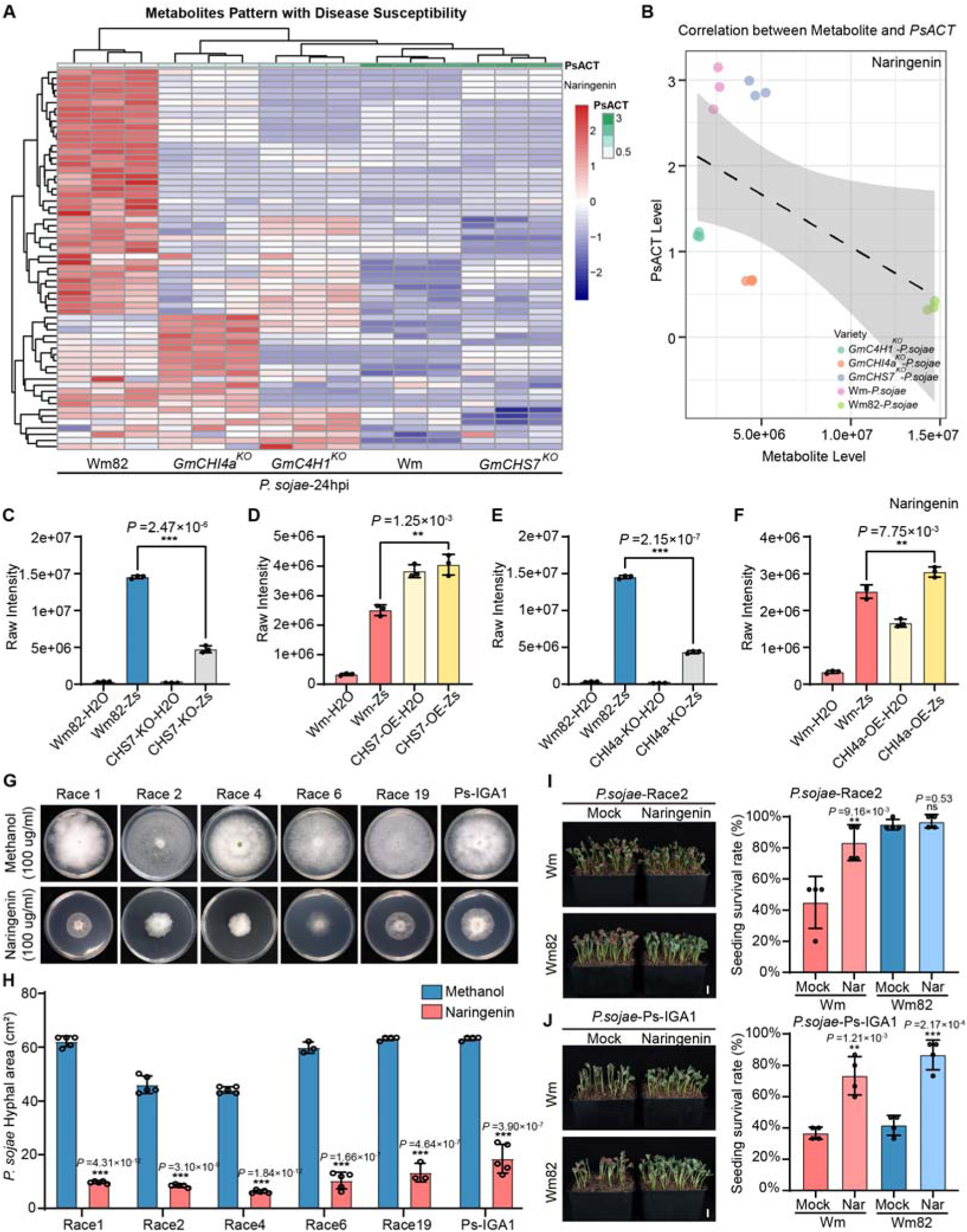
The flavonoid phytoalexin naringenin confers broad-spectrum resistance to *P. sojae* in soybean. (A) Heatmap of metabolite abundance patterns across soybean genotypes and treatments, aligned with disease susceptibility (*PsACT*). (B) Correlation analysis between naringenin abundance and *PsACT* across genotypes, showing a negative association between naringenin levels and *P. sojae* activity. (C–F) Quantification of naringenin levels in roots of *GmCHS7^KO^*, *GmCHS7^OE^*, *GmCHI4a^KO^*, and *GmCHI4a^OE^* lines at 24 hours post-inoculation with *P. sojae*. Knockout lines exhibit reduced naringenin accumulation, while overexpression lines show elevated levels, correlating with increased pathogen resistance. (G) Growth of multiple *P. sojae* races on medium supplemented with naringenin or methanol control. Naringenin strongly inhibits hyphal growth across various strains. (H) Quantification of hyphal growth area for each *P. sojae* race in the presence or absence of naringenin, demonstrating significant growth inhibition by naringenin across diverse pathotypes. (I, J) Survival of soil-grown Wm (susceptible) and Wm82 (resistant) seedlings pretreated with naringenin via seed coating and inoculated with *P. sojae* races R2 (I) or Ps-IGA1 (J). Naringenin significantly increases survival in Wm and further enhances resistance to Ps-IGA1 in Wm82. n ≥ 60 seedlings per treatment. Scale bar, 2 cm.

Naringenin, a product of CHS and type I/II CHI (Winkel-Shirley, 2001; Ralston et al., 2005), showed a significant negative correlation with susceptibility to *P. sojae*, with Pearson correlation coefficients -0.52 (*p*=0.04) in Wm82, the Wm82 background CRISPR mutants and Wm (Figure 6B). Meanwhile, the contents of naringenin were constitutively elevated by the overexpression of CHS7 and CHI4 (Figure 6D,F). To test if naringenin has a direct function in inhibiting *P. sojae* growth in host plants, we conducted a growth assay to assess the effects of naringenin on the hyphal growth of *P. sojae* Race 2 cultured on 10% V8 medium. *P. sojae* was inoculated onto media supplemented with various concentrations of the metabolites and incubated at 28°C for 7 days. The results demonstrated that naringenin exhibited dose-dependent inhibitory effects on Race 2 hyphal growth (Figure S8).

We next examined whether naringenin exerts similar effects on other *P. sojae* races. As shown in Figure 6G, the metabolites not only inhibited the growth of the R2 strain but also suppressed the growth of five other major pathogenic strains, including Ps-IGA1—a hypervirulent strain we recently identified from the field that overcomes *Rps1k*-triggered immunity. The virulence patterns of the tested strains is showed in Table S8.

Furthermore, we evaluated whether exogenous naringenin application could protect soybean plants from *P. sojae*–induced root rot under soil conditions. As shown in Figure 6I, seed coating with naringenin markedly enhanced plant survival: for the susceptible cultivar Wm, survival increased from 45% to 83.33% upon infection with R2, and from 36% to 73% with Ps-IGA1. In the resistant cultivar Wm82, naringenin treatment did not alter survival under R2 infection, consistent with its complete resistance to this pathotype. However, under Ps-IGA1 infection, naringenin application elevated the survival rate of Wm82 from approximately 41% to 86.6%. Collectively, these results demonstrate that naringenin effectively suppresses the growth and pathogenicity of diverse *P. sojae* isolates both in vitro and in soil. The incorporation of naringenin into seed coatings therefore holds considerable potential for protecting soybean against *P. sojae* root rot.

### 7. Spatial metabolic analysis of Wm and Wm82 under *P. sojae* infection reveals tissue-specific accumulation of flavonoid phytoalexins

Phenolic compounds, including flavonoids and their precursors, are known to emit bright blue autofluorescence, often associated with resistance to pathogens and pests (Chappell and Hahlbrock 1984; Hutzler, 1998). Recently, researchers documented a pattern switch of this blue autofluorescence before and after insect infection (Materska et al., 2022). Here using confocal microscopy, we observed that under control conditions, bright blue autofluorescence was evenly distributed in root cross-sections of both Wm and Wm82 (Figure 7A). However, following infection, the autofluorescence significantly diminished in Wm, whereas in Wm82, a distinct bright blue autofluorescence band surrounding the stele appeared under UV excitation in response to *P. sojae* infection (Figure 7A, Figure S9A,B).

**Figure 7.**
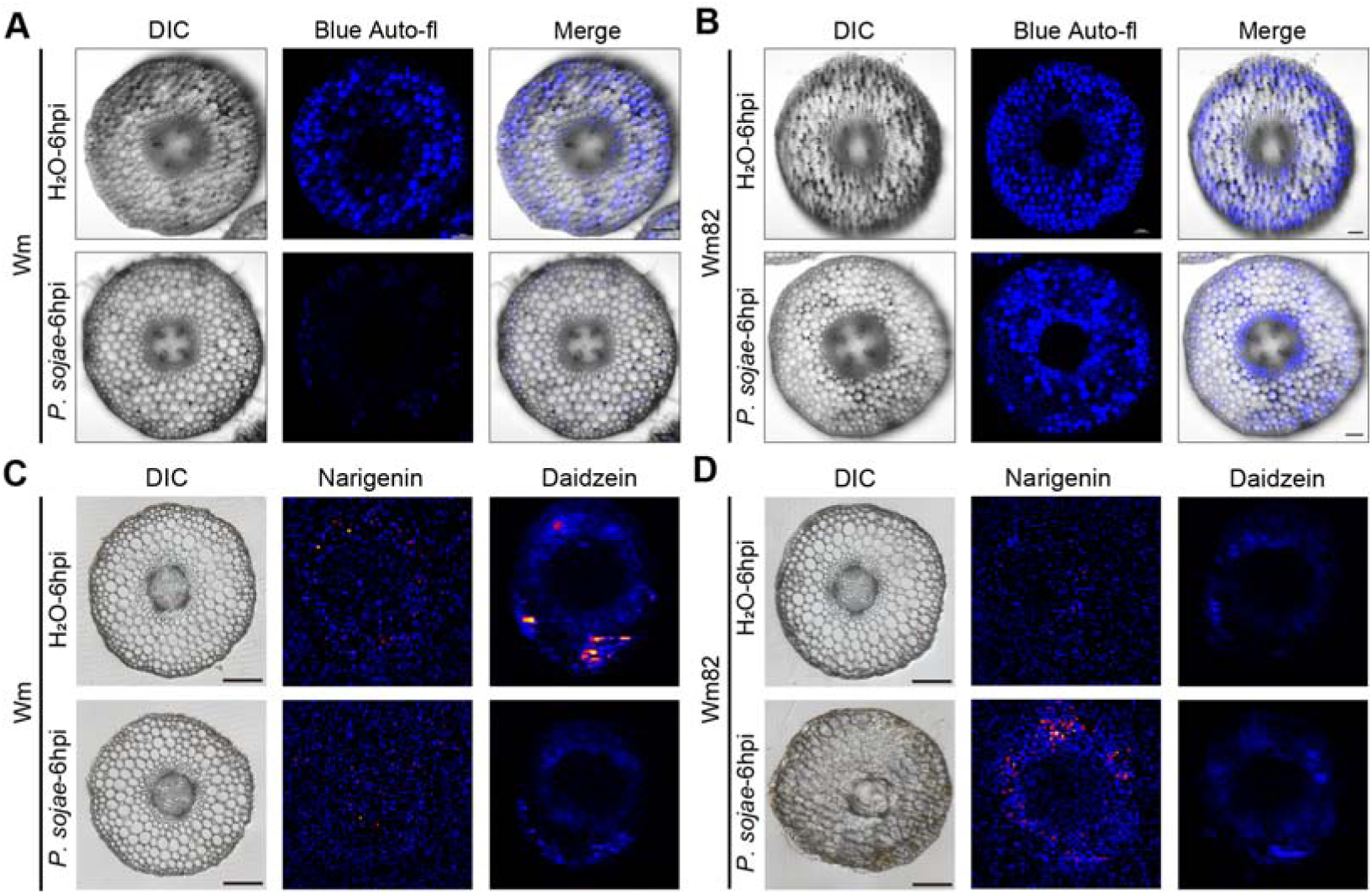
Flavonoid phytoalexin accumulation in response to *P. sojae* infection in Soybean roots. (A) Confocal microscopy images of root cross-sections from the susceptible cultivar Wm at 6 hours post-inoculation (hpi) with *P. sojae*. In control conditions (H_2_O), blue autofluorescence is evenly distributed in both Wm. Following infection, autofluorescence significantly diminishes in Wm. (B) Confocal microscopy images of root cross-sections from the resistant cultivar Wm82 at 6 hours post-inoculation (hpi) with *P. sojae*. In control conditions (H_2_O), blue autofluorescence is evenly distributed in Wm82. Following infection, autofluorescence exhibits a distinct autofluorescent band surrounding the stele, indicative of a localized immune response. (C) Spatial metabolic analysis of Wm roots at 6 hpi with *P. sojae*. Naringenin is bearly detectable in Wm. Daidzein was detected under control, while not detectable when infected by *P. sojae*. (D) Spatial metabolic analysis of Wm82 roots at 6 hpi, showing that naringenin forms a pronounced ring-like enrichment in stele-adjacent tissues. Daidzein was not detected either before or after infection. Scale bar, 100 μm

This observation recalled that the flavonoid synthesis genes were enriched in the pericycle of *P. sojae* infected Wm82. We hypothesize that this infection-triggered bright blue autofluorescence relocation corresponds to pericycle-enriched flavonoids synthesis in response to *P. sojae* infection. To validate this, we analyzed a CRISPR knockout of *GmCHS7* and *GmCHI4A* in the Wm82 genetic background. In these knockout samples, the autofluorescence band was absent following *P. sojae* infection (Figure S9C,D). Whereas in 35s promoter driven *GmCHS7* and *GmCHI4A* overexpression lines under Wm background, the blue autofluorescence showed ubiquitous distribution under both control condition and after infection (Figure S9E,F). These results were strongly correlated with the metabolic analysis results, providing strong evidence of a link between a functional flavonoid biosynthesis pathway, the formation of the pericycle-enriched autofluorescence band and resistance against *P. sojae*.

We speculate that flavonoids, especially naringenin, accumulate in tissues surrounding the stele, such as the pericycle, preventing the pathogen from invading the central cylinder, thereby conferring higher resistance in Wm82. To test our hypothesis, we performed a spatial metabolic analysis of Wm and Wm82 at 6 hpi with *P. sojae* (Figure 7). As shown in Figure 7D, naringenin was significantly enriched in tissues surrounding the stele specifically under Wm82 background in response to *P. sojae* infection. In contrast, under Wm background, we did not observe any increase in terms of naringenin contents at 6 hpi (Figure 7C). We also examined the spatial distributions of daidzein, which was previously demonstrated to facilitate infection (Graham, 1991; V. V. Lozovaya, 2004), under both Wm and Wm82 background. As shown in Figure 7C, daidzein was detected in roots in absence of infection and it was no longer detected at 6 hpi. In contrast, in Wm82, daidzein was not detected either before or after infection (Figure 7D).

These results highlight the critical role of tissue-specific phytoalexin accumulation in resistance to *P. sojae*. We propose that these phytoalexins contribute to the formation of a chemical barrier in tissues surrounding the stele, such as the pericycle, effectively preventing the pathogens from invading the stele and compromising vascular integrity.

## Discussion

In this study, we establish a comprehensive mechanistic framework for soybean resistance to *P. sojae*. By integrating pathogen imaging, single-cell transcriptomics, genetic manipulation, metabolic profiling, and spatial metabolite imaging, we demonstrate that resistance in the cultivar Williams 82 (Wm82), which carries *Rps1k*, is mediated by pericycle-enriched activation of flavonoid biosynthesis. While the pathogen initially colonizes the outer cortical tissues in both resistant and susceptible genotypes, its progression into the stele is specifically blocked in Wm82. This spatial restriction coincides with the emergence of an immune hotspot in the pericycle, marked by transcriptional reprogramming of flavonoid biosynthetic genes and accumulation of phytoalexins such as naringenin. Functional analyses using CRISPR-Cas9 knockouts and overexpression lines confirm that *GmCHS7* and *GmCHI4A* are essential for resistance. Moreover, naringenin not only inhibits *P. sojae* growth in vitro but also confers broad-spectrum protection in planta. Collectively, our findings uncover a previously unrecognized tissue-specific immune strategy in which flavonoid-derived metabolic barriers prevent vascular colonization and ensure resistance to a destructive oomycete pathogen.

### Spatial organization of plant immunity

These results directly address a central question in plant immunity: how immune responses are spatially organized across tissues and cell types to confer effective defense. Classical models often treat plant tissues as immunologically uniform, yet accumulating evidence indicates that cell-type-specific immunity plays a decisive role in restricting pathogen spread. In *Arabidopsis thaliana*, for instance, spatial expression of pattern recognition receptors (PRRs) and immune signaling components in epidermal tissues has been well characterized (Zhou et al., 2020). Recent work further shows that a pericycle-localized calcium-permeable cation channel mediates resistance to *Plasmodiophora brassicae* by activating tissue-specific defense (Wang et al., 2023). Similar tissue localization has been observed in *Brassica napus*, where stele-associated cells induce defense genes following infection by *Verticillium longisporum* (Johansson et al., 2006), and in tomato and *Arabidopsis*, where xylem parenchyma cells contribute to resistance against vascular wilt pathogens (Yadeta and Thomma, 2013). However, direct mechanistic evidence linking pericycle-specific transcriptional states to immune outcomes has been lacking. Our findings extend this framework by showing that flavonoid biosynthesis in the pericycle establishes a chemical barrier that complements structural defenses, thereby excluding *P. sojae* from vascular tissues. The single-cell atlas generated here demonstrates that immune activation is not uniformly distributed but is strongly enriched in the pericycle during incompatible interactions, offering a molecular explanation for the long-observed histological phenomenon of pathogen exclusion from the stele (Enkerli et al., 1997). By connecting pathogen invasion dynamics with transcriptional and metabolic reprogramming, we identify the pericycle as a central immune barrier in root defense.

### Immune zoning and pericycle activation

Unlike earlier reports describing hypersensitive responses (HR) and cell wall appositions at ∼4 h post-inoculation (Enkerli et al., 1997), Wm82 exhibits delayed cell death, beginning around 24 hpi, with visible oospore formation by 12 hpi. Despite early cortical colonization, pathogen proliferation is confined to outer tissues, suggesting that immune “zoning” occurs within the root—where specific cell layers adopt specialized defensive roles. To test this, we conducted single-cell transcriptomic analysis on both resistant and susceptible cultivars under *P. sojae* infection. Rather than the commonly used single-nucleus approach, we generated root protoplasts to enable the detection of both host transcripts and potential RNA trafficking events. Previous research has shown that *Ustilago maydis* exports mRNA to the host to manipulate its immune response (Kwon et al., 2021). To detect pathogen mRNA within host cells, reads were also mapped to a combined pseudo-genome of Wm82 and *P. sojae* (Espindula et al., 2020), facilitating simultaneous host–pathogen transcriptomic profiling (Table S9). Comparative analyses across cell types may elucidate how host physiology influences pathogen gene expression and RNA mobility. However, further validation is still needed to distinguish whether the pathogen reads originated from broken hyphae in the environment or were truly internalized within host cells,

The resulting atlas comprises 15 transcriptionally distinct cell clusters. Among them, cluster C2, annotated as pericycle, exhibited the highest transcriptional activity, with over 70% of Wm82-specific upregulated genes during infection localized to this cluster. Pseudotime trajectory analysis revealed a pathogen-induced transition toward an immune-activated pericycle state enriched for genes associated with systemic acquired resistance, ATP generation, and specialized metabolite biosynthesis (Table S5). Thus, the pericycle acts not merely as a structural boundary but as a metabolically dynamic immune hub.

### Naringenin as a central phytoalexin and metabolic marker

Among the identified metabolites, naringenin emerged as a functional phytoalexin showing a strong inverse correlation with pathogen biomass across genotypes. Its accumulation was strictly dependent on *CHS7* and *CHI4A*, and in vitro assays confirmed its dose-dependent inhibition of *P. sojae*. Remarkably, exogenous naringenin also conferred resistance to race Ps-IGA1, which circumvents *Rps1k*-mediated recognition. While isoflavonoids such as glyceollins have long been considered major defensive compounds in legumes (Graham, 1991), our results highlight flavanones like naringenin as equally potent and more broadly effective antimicrobial metabolites. Thus, naringenin is both necessary and sufficient to restrict pathogen spread, even when classical *R*-gene-mediated immunity fails.

### Flavonoid autofluorescence and visualization of immune zoning

A pericycle-localized ring of blue autofluorescence emerged 24 h after *P. sojae* infection in a *CHS7*-dependent manner. This pattern closely matched naringenin accumulation and correlated strongly with resistance intensity. Although autofluorescence spectra can vary with excitation conditions and microenvironmental factors, the observed blue signal in unstained root cross-sections coincided spatially with flavonoid deposition and was absent in the susceptible genotype. Combined with genetic evidence showing loss of autofluorescence in flavonoid-deficient lines, these results indicate that the blue signal primarily reflects defensive flavonoid deposition at the pericycle–cortex interface, with a possible minor contribution from phenolic polymers in the cell wall.

### Conceptual advances and implications

Together, these results advance the concept of *spatial immunity*—the idea that the efficacy of defense depends on its cellular and tissue-specific deployment. The “immune hotspot” model proposed here integrates transcriptional and metabolic evidence to illustrate that defense activation in Wm82 is spatially and temporally targeted to protect vascular integrity (Chuberre et al., 2018). Rather than broad immune activation, the resistant cultivar executes a focused, energy-efficient response.

### Limitations and future directions

Despite these insights, several key questions remain. Our analyses primarily capture early infection events, whereas later stages involving vascular invasion remain unexplored due to extensive cell death that limits protoplast recovery. Implementing single-nucleus RNA-seq across multiple time points could overcome this limitation by providing a temporal trajectory of both host and pathogen transcriptomes. Moreover, although we identify flavonoid biosynthesis as a critical downstream defense pathway, the upstream signaling cascade linking *Rps* recognition to pericycle activation is still undefined. Potential mediators may include transcription factors, kinases, or hormone-responsive regulators that warrant further investigation. Addressing these questions will deepen our understanding of spatially resolved immunity and inform the development of precision-based strategies for durable crop protection.

### Applications for crop improvement

Our findings suggest practical avenues for engineering durable resistance. The flavonoid biosynthetic pathway—particularly *CHS7* and *CHI4A*—offers promising molecular targets. Breeding programs could prioritize alleles or regulatory elements driving pericycle-specific expression, or employ cell-type-specific promoters to boost flavonoid accumulation precisely where it is most effective. Furthermore, the identification of naringenin as a potent phytoalexin supports its potential use as an exogenous defense elicitor during seedling establishment or under high disease pressure, complementing genetic resistance.

Finally, the spatial precision of immune deployment observed in Wm82 implies a broader principle: targeted activation of immune-competent cell types can enhance disease resistance without incurring major growth penalties. Such *spatially optimized defense engineering*—focusing on metabolically active boundary tissues like the pericycle—may represent a next-generation paradigm for sustainable crop immunity.

## Methods

### Plant materials and growth conditions

The soybean cultivars Williams (Wm) and Williams 82 (Wm82) were obtained from Shandong Agricultural University. *Glycine max* L. cv. Wm82 and Wm seeds were surface-sterilized with chlorine gas (generated from 100 mL NaClO and 4 mL concentrated HCl). Plants were subsequently grown in a phytotron under a 16-h-light/8-h-dark photoperiod at 25 °C and 65% relative humidity.

### Pathogen Cultivation and Zoospore Production

The *P. sojae* strains Race 2, Race 1, Race 4, Race 6, Race 19 and Race 2-YFP obtained from Wang Lab in Shandong Agricultural University. Ps-IGA1 was isolated from field. The selected strains were inoculated on 10% V8 agar medium and incubated in the dark at 25 °C for 7 days. Mycelial plugs (8 mm diameter) were then transferred to 10% V8 juice medium and incubated at 25 °C for an additional 5 days in the dark. After incubation, mycelia were rinsed five times with sterilized ddH□O. Subsequently, 10 mL of sterilized ddH□O was added to each plate, and the plates were incubated at 25 °C for 8–10 hours. Zoospore suspension was collected once zoospores were released in large quantities.

### *P. sojae* Inoculation of Soybean Seedlings

Soybean seeds were sown in plastic pots containing vermiculite and incubated at 25 °C in the dark for 3 days. Zoospores were prepared as described previously and diluted to 1 × 10³ zoospores/mL. Etiolated seedling roots were inoculated by pipetting 5 mL of zoospore suspension onto the roots. The plants were maintained in a climate-controlled room at 25 °C and 80% relative humidity, kept in the dark. Roots were harvested at 4 hours post-inoculation (hpi).

### Sample preparation and library construction for single cell RNA sequencing

We employed a modified version of the protoplast isolation method originally described by Jia et al. Whole roots were harvested from each replicate three days post-sowing, precisely four hours after treatment with either distilled water or *P. sojae* Race 2 zoospores. The roots were first cleared of soil debris by rinsing with tap water, then immersed in plasmolysis buffer in a 60-mm Petri dish. The roots were subsequently sliced into 0.5–1 mm segments using a surgical blade. To remove slicing-induced debris, the root segments were pre-washed in a 70-μm cell strainer with plasmolysis buffer. The segments were then weighed, and each sample was transferred to 20 mL of digestion buffer in separate, clean 60-mm Petri dishes. A 30-min vacuum infiltration at 0.7 kg/cm² was performed to accelerate plasmolysis.

Samples were incubated for 4 h at room temperature with gentle shaking at 40 rpm in the dark. After digestion, the protoplast-enzyme mixture was cooled on ice, filtered sequentially through 70-μm and 40-μm cell strainers, and collected in 50-mL conical tubes. Protoplasts were washed with 10 mL of pre-cooled W5 buffer, centrifuged at 400 × *g* for 2 min at 4 °C, and resuspended in 300 μL of plasmolysis buffer. Cell counts were performed using a hemocytometer, and the density was adjusted to ∼1,000 cells/μL. For single-cell sequencing, ∼20,000 cells from each replicate were loaded onto a 10x Genomics Chromium Chip A using v3 chemistry, targeting the capture of 5,000 cells. Subsequent steps, including cell barcoding, reverse transcription, and library construction, followed the manufacturer’s protocols. Sequencing was performed by Novogene Co., Ltd

### Single cell RNA seq data analysis

Sequencing reads were processed and aligned to the *Glycine max* v4.0 reference genome (NCBI) using the Cell Ranger (v4.0, 10× Genomics) pipeline with default settings. Downstream analyses were performed in Seurat v5.0.3 (Butler et al., 2018). Cells were retained if they expressed 200–6,000 genes and contained 500–20,000 unique molecular identifiers (UMIs). Data from individual samples were integrated using the *FindIntegrationAnchors* and *IntegrateData* functions in Seurat. Principal component analysis (PCA) was used for dimensionality reduction, and cell clusters were visualized using Uniform Manifold Approximation and Projection (UMAP) at a resolution of 1.0 based on the top 20 PCs.

Cluster-specific marker genes were identified using the COSG package (Wu et al., 2025). Arabidopsis homologs of COSG markers were inferred using the *OthoFinder* R interface (Emms and Kelly, 2019). These homologs were cross-referenced against the marker gene library of Shahan et al. COSG markers whose homologs appeared in the Shahan atlas were retained for annotation. In addition, COSG markers were cross-checked against published soybean root marker sets (Miao and Verma, 1993; Nontachaiyapoom et al., 2007; Wang et al., 2022; Yan et al., 2022; Sun et al., 2023; Zhang et al., 2025). Markers exhibiting clear cluster-specific expression were assembled into a marker table for SCINA-based cell-type annotation (Zhang et al., 2019). Average expression levels and the proportion of marker-expressing cells were visualized using bubble plots (Figure S5).

Differential gene expression (DGE) analyses were performed using *FindMarkers* in Seurat with thresholds of |log2 fold change| ≥ 0.25 and adjusted *P* ≤ 0.05. UCell scores for selected gene sets were computed and visualized using the UCell package (Andreatta and Carmona, 2021) and scCustomize (https://samuel-marsh.github.io/scCustomize/). Pseudotime trajectories were reconstructed using Monocle 2 (Trapnell et al., 2014; Qiu et al., 2017). Co-expression modules were generated using hdWGCNA (Morabito et al., 2023), and modules specifically activated in Wm82 during infection were used to build a gene regulatory network, which was visualized in Cytoscape (Shannon et al., 2003).

### Microscopy

At 4, 6, 12, 24, and 48 hpi, the roots of Wm and Wm82 seedlings inoculated with *P. sojae* strain Race2-YFP were cut near the meristem and subjected to the preparation of 5 μm-thick transverse slices by free-hand sectioning. The slices were observed under a laser confocal microscope (C2, Nikon, Japan) to monitor the growth of *P. sojae* hyphae. YFP fluorescence was visualized using excitation and emission wavelengths of 488 and 500–550 nm, respectively.

### In situ hybridization

Soybean seedlings cultivated in the dark at 3 days were fixed in FAA solution for 24 h at 4 °C and dehydrated in an ethanol series, cleared in a xylene series and embedded in paraffin. Then, 10 µm sections were prepared using a microtome (RM2245, Leica, Germany). For probe labelling, specific 500 bp sequences of target genes were amplified from the complementary DNA (cDNA) of Williams 82 with primers and cloned to pGEM®-T Easy Vector (Promega, USA, A1360). With the resulting vectors as templates, both SP6 and T7 promoter-fused fragments were amplified and purified. Then the digoxigenin-labelled antisense and sense probes were in vitro transcribed with these fragments as templates, respectively using SP6 and T7 RNA polymerase with a DIG RNA labelling kit (Roche, Switzerland, 11175025910). The paraffin on the sections was dissolved with xylene, and the tissue was digested with Proteinase K (Sigma, USA, P2308) and dehydrated with graded alcohol. The prepared probe buffer was added to the tissue sections and incubated for 12 h at 55 °C in a humidified box. After hybridization, the sections were incubated with anti-digoxin antibody (Roche, Switzerland, 11093274910) for 90 min. Then the sections were washed and incubated in a colour reaction solution (Roche, Switzerland, 11681451001) containing blue tetrazolium chloride and 5-bromo-4-chloro-3-indolylphosphate for 36 h in a dark humidified box. Photographs were taken under bright-field illumination using a microscope (DM2500, Leica, Germany). For each probe, at least three roots were analysed, and all showed the same expression pattern.

### Plasmid construction

First, the CDS sequence of gene was analysed using the CCTop - CRISPR/Cas9 target online predictor (https://cctop.cos.uni-heidelberg.de:8043). A pair single-guide RNAs (sgRNAs) highly specific oligonucleotides to the gene of interest were annealed and cloned into a *PGEs201* knockout expression vector. The full-length gene sequences were amplified from Wm82 using overlapping PCR. The PCR amplicons and eGFP sequence were inserted into the binary vector *pCAMBIA3301H* between the restriction endonuclease sites SacI and XbaI to generate plasmid. The plasmids were then transformed into *Agrobacterium* strain EHA105 and K599 for transformation.

### Stable and Transient Transformation

*Agrobacterium rhizogenes*-mediated hairy root transformation was conducted following the method described by Fan et al. (Fan et al., 2020) with minor modifications. Soybean seeds were surface-sterilized using chlorine gas and germinated in a phytotron for 4 days. Agrobacterium cultures were prepared and inoculated at the cotyledonary node and/or the hypocotyl region proximal to the cotyledons. After inoculation, the seedlings were transferred to moist sterile vermiculite and maintained in a growth chamber for 2–3 weeks to allow hairy root induction. When the emerging hairy roots reached approximately 5–10 cm in length, the transgenic roots carrying the resistance construct were identified and selected.

A slightly modified soybean transformation protocol previously described by Yamada et al. (Yamada et al., 2010) was employed. Sterilized soybean seeds were germinated on B5 medium overnight, and the embryos with cotyledons were excised by removing the shoot apices and seed coats. The explants were wounded with a scalpel and immersed in the *Agrobacterium* infection solution. After five days of co-cultivation, explants were transferred to shoot-induction medium for initial regeneration, followed by selection on medium containing 10 mg mL□¹ glufosinate to induce cluster-shoot formation. The regenerated shoots were then moved to elongation medium with glufosinate until they reached approximately 5 cm in length. Elongated shoots were transferred to rooting medium, and rooted plantlets (3–5 cm root length) were transplanted into soil and grown in a controlled growth chamber (14 h light/10 h dark, 26 °C).

### qPCR analysis

Total RNA was extracted from soybean roots with TRIzol (Tiangen, China, DP424). RNA concentration and quality were determined using a NanoDrop spectrophotometer (N50; Implen, Germany). First□strand cDNA was synthesized from 1 μg of RNA using the StarScript III All-in-one RT Mix with gDNA Remover (GenStar, China, A236) according to the manufacturer’s instructions. The cDNA was diluted to 100 ng/μl in sterile water, and 1 μl diluent was used as the template for RT-qPCR. RT-qPCR was performed using 2×RealStar Fast Mixture (GenStar, China, A301) on a QuantStudio 1 Sequence Detection System (Q1, Applied Biosystems, USA) following the manufacturer’s instructions. Expression levels were quantified using three biological replicates. The differences between groups were calculated with the 2^-ΔΔCt^ method. Relative expression levels were calculated using *Actin11* (*Glyma.18G290800*) as an internal control. The primers used for RT-qPCR are listed in Table S10.

### *P. sojae* Biomass Assays

Transgenic soybean plants were inoculated with *P. sojae* strain Race 2. Each sample contains roots from 3 individual *P. sojae* infected plants. DNA was extracted using a plant DNA kit (Tiangen, China, DP350) for the biomass assay. The virulence was quantified by determining the ratio between *P. sojae* DNA to soybean DNA in infected plants using quantitative PCR. All assays were repeated independently at least three times. Primer sequences used for PCR are listed in Table S10.

### LC-MS/MS Sample Preparation and Analysis

Samples were freeze-dried and ground into powder. For extraction, 0.04 g of powder was mixed with 70% methanol containing an internal standard. The mixture was vortexed for 15 minutes at room temperature, incubated at -20 °C for 30 minutes, and centrifuged at 12,000 rpm for 3 minutes at 4 °C. The supernatant was filtered through a 0.22 μm PTFE membrane and stored at -20 °C until analysis.

LC-MS/MS analysis was performed using a UPLC-ESI-QTRAP system (ExionLC™ AD, Sciex, USA) with an Agilent SB-C18 column (1.8 μm, 2.1 × 100 mm). The mobile phase consisted of water with 0.1% formic acid (A) and acetonitrile with 0.1% formic acid (B), using a linear gradient from 95% A to 95% B. The flow rate was set to 0.35 mL/min, the column temperature was maintained at 40 °C, and the injection volume was 2 μL. The ESI source was operated at 500 °C with an ion spray voltage of 5500 V (positive) or -4500 V (negative). Multiple reaction monitoring (MRM) mode was used for quantification, with optimized declustering potential and collision energy for each metabolite.

### DESI-MSI Analysis

Soybean root tissues were embedded in 7% agarose and sectioned transversely to 80 μm thickness using a vibrating microtome (VT1000S, Leica, Germany) (Zhang et al., 2022). The sections were transferred onto Superfrost Plus glass slides (Fisher Scientific, USA, 12-550-15) and vacuum-dried in a desiccator (Dong et al., 2020).

DESI Mass Spectrometry Imaging was performed using a SYNAPT XS mass spectrometer equipped with a DESI-XS ion source (Waters, United Kingdom). The spray solvent (methanol:water:formic acid, 98:2:0.1, v/v/v) was delivered at a flow rate of 3 μL/min. The capillary voltage was set to 0.5 kV, and nitrogen was used at 0.5 bar for nebulization. A heated transfer line at 255 °C was used for desolvation and ion transmission. The pixel size was set to 20 μm, with a grating speed of 400 μm/s during acquisition. Mass spectra were collected in positive-ion full-scan mode, with a sampling cone voltage of 40 V, over an m/z range of 50–1500 Da.

Data were collected using MassLynx Software V4.2 (Waters Corporation). HDI v1.6 software was used to export data into imzML format (Waters, United Kingdom), which was then imported into ShinyCardinal V0.3.5 (Bemis et al., 2023; Dong and Heining, 2024) for data preprocessing and visualization.

### Seed Coating Formulation and Application

The seed-coating film solution was prepared by dissolving polyvinyl alcohol (PVA) in water to a final concentration of 5% (w/v) and stirring at room temperature overnight. Gum arabic (0.5–1% w/v) was added to the PVA solution to enhance coating adhesion. Naringenin was incorporated into the coating material by adding it to the film-forming solution at a final concentration of 100 μg/mL, followed by thorough mixing to achieve uniform dispersion. Talc powder (1–2% w/v) was added as a filler to adjust viscosity, and food-grade dye was included for color labeling.

For seed coating, the mixture was applied to the seeds at a ratio of 5–10% (v/w, coating solution: seeds) while mixing continuously to ensure even coverage. After coating, the seeds were air-dried at room temperature until the coating layer solidified. The coated seeds were then inoculated with *P. sojae* for disease-resistance assays.

### Statistical Analysis

Phenotypic analysis and figure generation for *P. sojae* biomass and hyphal area were performed using GraphPad Prism (version 9.0). YFP fluorescence signals corresponding to hyphal invasion tracks were quantified using ImageJ. Fluorescence intensity was measured within manually defined regions of interest, and the extracted values were used for subsequent statistical analyses. Statistical comparisons were performed using two-tailed Student’s t-tests or one-way ANOVA, as appropriate. Statistical significance was defined as follows: *p* < 0.05 (*), *p* < 0.01 (**), and *p* < 0.001 (***).

## ACKNOWLEDGMENTS

We sincerely thank Prof. Yuefeng Guan (School of life sciences Guangzhou University) for providing the PGES201 vector used for CRISPR and Prof. Fanli Meng (Northeast Institute of Geography and Agroecology, CAS) for providing the RUBY construct used for hairy-root transformation. This work was supported by the Agriculture Science and Technology Major Project and the Chinese Academy of Science (E329S502).

## AUTHOR CONTRIBUTIONS

Y.G., Q.Q.W., and J.Z. conceived and designed the experiments. Y.G. prepared ScRNAseq samples, Y.G., Q.S.W., and H.W. performed data analysis. Q.S.W., Z.D., R.Z., C.M., and X.W. performed disease assay and plant gene function analysis. Z.D. and Q.Q.W. contributed soybean materials and *Phytophthora sojae* to this study. Y.G., Q.Q.W. and J.Z. wrote the manuscript.

## Declaration of generative AI and AI-assisted technologies in the manuscript preparation process

During the preparation of this work the authors used ChatGPT and deepseek to improve writing (only for text polishing and grammar correction). After using this tool, the authors reviewed and edited the content as needed and take full responsibility for the content of the published article.

## References

1. Andreatta, M., and Carmona, S.J. (2021). UCell: Robust and scalable single-cell gene signature scoring. Comput Struct Biotechnol J 19, 3796–3798.

2. Bemis, K.A., Foll, M.C., Guo, D., Lakkimsetty, S.S., and Vitek, O. (2023). Cardinal v.3: a versatile open-source software for mass spectrometry imaging analysis. Nat Methods 20, 1883–1886.

3. Bradley, C.A., Allen, T.W., Sisson, A.J., Bergstrom, G.C., Bissonnette, K.M., Bond, J., Byamukama, E., Chilvers, M.I., Collins, A.A., Damicone, J.P., Dorrance, A.E., Dufault, N.S., Esker, P.D., Faske, T.R., Fiorellino, N.M., Giesler, L.J., Hartman, G.L., Hollier, C.A., Isakeit, T., Jackson-Ziems, T.A., Jardine, D.J., Kelly, H.M., Kemerait, R.C., Kleczewski, N.M., Koehler, A.M., Kratochvil, R.J., Kurle, J.E., Malvick, D.K., Markell, S.G., Mathew, F.M., Mehl, H.L., Mehl, K.M., Mueller, D.S., Mueller, J.D., Nelson, B.D., Overstreet, C., Padgett, G.B., Price, P.P., Sikora, E.J., Small, I., Smith, D.L., Spurlock, T.N., Tande, C.A., Telenko, D.E.P., Tenuta, A.U., Thiessen, L.D., Warner, F., Wiebold, W.J., and Wise, K.A. (2021). Soybean Yield Loss Estimates Due to Diseases in the United States and Ontario, Canada, from 2015 to 2019. Plant Health Progress 22, 483–495.

4. Butler, A., Hoffman, P., Smibert, P., Papalexi, E., and Satija, R. (2018). Integrating single-cell transcriptomic data across different conditions, technologies, and species. Nat Biotechnol 36, 411–420.

5. Chappell, J., and Hahlbrock, K. (1984). Transcription of plant defence genes in response to UV light or fungal elicitor. Nature 311, 3.

6. Chuberre, C., Plancot, B., Driouich, A., Moore, J.P., Bardor, M., Gugi, B., and Vicre, M. (2018). Plant Immunity Is Compartmentalized and Specialized in Roots. Front Plant Sci 9, 1692.

7. Dong, Y., and Heining, U. (2024). Mass Spectrometry Imaging Data Analysis with ShinyCardinal. Research Square.

8. Dong, Y., Sonawane, P., Cohen, H., Polturak, G., Feldberg, L., Avivi, S.H., Rogachev, I., and Aharoni, A. (2020). High mass resolution, spatial metabolite mapping enhances the current plant gene and pathway discovery toolbox. New Phytol 228, 1986–2002.

9. Emms, D.M., and Kelly, S. (2019). OrthoFinder: phylogenetic orthology inference for comparative genomics. Genome Biol 20, 238.

10. Enkerli, K., Mims, C.W., and Hahn, M.G. (1997). Ultrastructure of compatible and incompatible interactions of soybean roots infected with the plant pathogenic oomycete Phytophthora sojae. Canadian Journal of Botany 75, 1493–1508.

11. Falcone Ferreyra, M.L., Rius, S.P., and Casati, P. (2012). Flavonoids: biosynthesis, biological functions, and biotechnological applications. Front Plant Sci 3, 222.

12. Fan, Y.L., Zhang, X.H., Zhong, L.J., Wang, X.Y., Jin, L.S., and Lyu, S.H. (2020). One-step generation of composite soybean plants with transgenic roots by Agrobacterium rhizogenes-mediated transformation. BMC Plant Biol 20, 208.

13. Gao, H., Narayanan, N.N., Ellison, L., and Bhattacharyya, M.K. (2005). Two classes of highly similar coiled coil-nucleotide binding-leucine rich repeat genes isolated from the Rps1-k locus encode Phytophthora resistance in soybean. Mol Plant Microbe Interact 18, 1035–1045.

14. Graham, T.L. (1991). Flavonoid and isoflavonoid distribution in developing soybean seedling tissues and in seed and root exudates. Plant Physiol 95, 594–603.

15. Hale, B., Brown, E., and Wijeratne, A. (2023). An updated assessment of the soybean–Phytophthora sojae pathosystem. Plant Pathology 72, 843–860.

16. Hutzler, P. (1998). Tissue localization of phenolic compounds in plants by confocal laser scanning microscopy. Journal of Experimental Botany 49, 953–965.

17. Johansson, A., Staal, J., and Dixelius, C. (2006). Early responses in the Arabidopsis-Verticillium longisporum pathosystem are dependent on NDR1, JA- and ET-associated signals via cytosolic NPR1 and RFO1. Mol Plant Microbe Interact 19, 958–969.

18. Kereszt, A., Li, D., Indrasumunar, A., Nguyen, C.D., Nontachaiyapoom, S., Kinkema, M., and Gresshoff, P.M. (2007). Agrobacterium rhizogenes-mediated transformation of soybean to study root biology. Nat Protoc 2, 948–952.

19. Lin, F., Chhapekar, S.S., Vieira, C.C., Da Silva, M.P., Rojas, A., Lee, D., Liu, N., Pardo, E.M., Lee, Y.C., Dong, Z., Pinheiro, J.B., Ploper, L.D., Rupe, J., Chen, P., Wang, D., and Nguyen, H.T. (2022). Breeding for disease resistance in soybean: a global perspective. Theor Appl Genet 135, 3773–3872.

20. Materska, M., Pabich, M., Sachadyn-Krol, M., Konarska, A., Weryszko-Chmielewska, E., Chilczuk, B., Staszowska-Karkut, M., Jackowska, I., and Dmitruk, M. (2022). The Secondary Metabolites Profile in Horse Chestnut Leaves Infested with Horse-Chestnut Leaf Miner. Molecules 27.

21. McCoy, A.G., Belanger, R.R., Bradley, C.A., Cerritos-Garcia, D.G., Garnica, V.C., Giesler, L.J., Grijalba, P.E., Guillin, E., Henriquez, M.A., Kim, Y.M., Malvick, D.K., Matthiesen, R.L., Mideros, S.X., Noel, Z.A., Robertson, A.E., Roth, M.G., Schmidt, C.L., Smith, D.L., Sparks, A.H., Telenko, D.E.P., Tremblay, V., Wally, O., and Chilvers, M.I. (2023). A global-temporal analysis on Phytophthora sojae resistance-gene efficacy. Nat Commun 14, 6043.

22. Miao, G.H., and Verma, D.P. (1993). Soybean nodulin-26 gene encoding a channel protein is expressed only in the infected cells of nodules and is regulated differently in roots of homologous and heterologous plants. Plant Cell 5, 781–794.

23. Mierziak, J., Kostyn, K., and Kulma, A. (2014). Flavonoids as important molecules of plant interactions with the environment. Molecules 19, 16240–16265.

24. Morabito, S., Reese, F., Rahimzadeh, N., Miyoshi, E., and Swarup, V. (2023). hdWGCNA identifies co-expression networks in high-dimensional transcriptomics data. Cell Rep Methods 3, 100498.

25. Morris, P.F., Bone, E., and Tyler, B.M. (1998). Chemotropic and contact responses of phytophthora sojae hyphae to soybean isoflavonoids and artificial substrates. Plant Physiol 117, 1171–1178.

26. Nontachaiyapoom, S., Scott, P.T., Men, A.E., Kinkema, M., Schenk, P.M., and Gresshoff, P.M. (2007). Promoters of orthologous Glycine max and Lotus japonicus nodulation autoregulation genes interchangeably drive phloem-specific expression in transgenic plants. Mol Plant Microbe Interact 20, 769–780.

27. Qiu, X., Hill, A., Packer, J., Lin, D., Ma, Y.A., and Trapnell, C. (2017). Single-cell mRNA quantification and differential analysis with Census. Nat Methods 14, 309–315.

28. Ralston, L., Subramanian, S., Matsuno, M., and Yu, O. (2005). Partial reconstruction of flavonoid and isoflavonoid biosynthesis in yeast using soybean type I and type II chalcone isomerases. Plant Physiol 137, 1375–1388.

29. Sahoo, D.K., Das, A., Huang, X., Cianzio, S., and Bhattacharyya, M.K. (2021). Tightly linked Rps12 and Rps13 genes provide broad-spectrum Phytophthora resistance in soybean. Sci Rep 11, 16907.

30. Schmitthenner, A.F. (1985). Problems and Progress in Control of Phytophthora Root Rot of Soybean. Plant Disease 69, 7.

31. Shahan, R., Hsu, C.W., Nolan, T.M., Cole, B.J., Taylor, I.W., Greenstreet, L., Zhang, S., Afanassiev, A., Vlot, A.H.C., Schiebinger, G., Benfey, P.N., and Ohler, U. (2022). A single-cell Arabidopsis root atlas reveals developmental trajectories in wild-type and cell identity mutants. Dev Cell 57, 543–560 e549.

32. Shannon, P., Markiel, A., Ozier, O., Baliga, N.S., Wang, J.T., Ramage, D., Amin, N., Schwikowski, B., and Ideker, T. (2003). Cytoscape: a software environment for integrated models of biomolecular interaction networks. Genome Res 13, 2498–2504.

33. Subramanian, S., Graham, M.Y., Yu, O., and Graham, T.L. (2005). RNA interference of soybean isoflavone synthase genes leads to silencing in tissues distal to the transformation site and to enhanced susceptibility to Phytophthora sojae. Plant Physiol 137, 1345–1353.

34. Sun, B., Wang, Y., Yang, Q., Gao, H., Niu, H., Li, Y., Ma, Q., Huan, Q., Qian, W., and Ren, B. (2023). A high-resolution transcriptomic atlas depicting nitrogen fixation and nodule development in soybean. J Integr Plant Biol 65, 1536–1552.

35. Trapnell, C., Cacchiarelli, D., Grimsby, J., Pokharel, P., Li, S., Morse, M., Lennon, N.J., Livak, K.J., Mikkelsen, T.S., and Rinn, J.L. (2014). The dynamics and regulators of cell fate decisions are revealed by pseudotemporal ordering of single cells. Nat Biotechnol 32, 381–386.

36. Treutter, D. (2005). Significance of flavonoids in plant resistance and enhancement of their biosynthesis. Plant Biol (Stuttg) 7, 581–591.

37. Tyler, B.M. (2007). Phytophthora sojae: root rot pathogen of soybean and model oomycete. Mol Plant Pathol 8, 1–8.

38. Tyler, B.M., Tripathy, S., Zhang, X., Dehal, P., Jiang, R.H., Aerts, A., Arredondo, F.D., Baxter, L., Bensasson, D., Beynon, J.L., Chapman, J., Damasceno, C.M., Dorrance, A.E., Dou, D., Dickerman, A.W., Dubchak, I.L., Garbelotto, M., Gijzen, M., Gordon, S.G., Govers, F., Grunwald, N.J., Huang, W., Ivors, K.L., Jones, R.W., Kamoun, S., Krampis, K., Lamour, K.H., Lee, M.K., McDonald, W.H., Medina, M., Meijer, H.J., Nordberg, E.K., Maclean, D.J., Ospina-Giraldo, M.D., Morris, P.F., Phuntumart, V., Putnam, N.H., Rash, S., Rose, J.K., Sakihama, Y., Salamov, A.A., Savidor, A., Scheuring, C.F., Smith, B.M., Sobral, B.W., Terry, A., Torto-Alalibo, T.A., Win, J., Xu, Z., Zhang, H., Grigoriev, I.V., Rokhsar, D.S., and Boore, J.L. (2006). Phytophthora genome sequences uncover evolutionary origins and mechanisms of pathogenesis. Science 313, 1261–1266.

39. V. V. Lozovaya, A.V.L., S. Li, G. L. Hartman, J. M. Widholm. (2004). Biochemical Response of Soybean Roots to Fusarium solani f. sp. glycines Infection. Crop Physiology & Metabolism 44, 8.

40. Wang, C., Li, M., Zhao, Y., Liang, N., Li, H., Li, P., Yang, L., Xu, M., Bian, X., Wang, M., Wu, S., Niu, X., Wang, M., Li, X., Sang, Y., Dong, W., Wang, E., Gallagher, K.L., and Wu, S. (2022). SHORT-ROOT paralogs mediate feedforward regulation of D-type cyclin to promote nodule formation in soybean. Proc Natl Acad Sci U S A 119.

41. Wang, W., Chen, L., Fengler, K., Bolar, J., Llaca, V., Wang, X., Clark, C.B., Fleury, T.J., Myrvold, J., Oneal, D., van Dyk, M.M., Hudson, A., Munkvold, J., Baumgarten, A., Thompson, J., Cai, G., Crasta, O., Aggarwal, R., and Ma, J. (2021). A giant NLR gene confers broad-spectrum resistance to Phytophthora sojae in soybean. Nat Commun 12, 6263.

42. Wang, W., Qin, L., Zhang, W., Tang, L., Zhang, C., Dong, X., Miao, P., Shen, M., Du, H., Cheng, H., Wang, K., Zhang, X., Su, M., Lu, H., Li, C., Gao, Q., Zhang, X., Huang, Y., Liang, C., Zhou, J.M., and Chen, Y.H. (2023). WeiTsing, a pericycle-expressed ion channel, safeguards the stele to confer clubroot resistance. Cell 186, 2656–2671 e2618.

43. Ward, E.W.B., Cahill, D.M., and Bhattacharyya, M.K. (1989). Early cytological differences between compatible and incompatible interactions of soybeans with Phytophthora megasperma f.sp. glycinea. Physiological and Molecular Plant Pathology 34, 17.

44. Winkel-Shirley, B. (2001). Flavonoid biosynthesis. A colorful model for genetics, biochemistry, cell biology, and biotechnology. Plant Physiol 126, 485–493.

45. Wu, S.J., Dai, M., Yang, S.P., McCann, C., Qiu, Y., Kumar, V., Marrero, G.J., Tsyporin, J., Huang, S., Shin, D., Stogsdill, J.A., Di Bella, D.J., Xu, Q., Chen, B., Farhi, S.L., Macosko, E.Z., Chen, F., and Fishell, G. (2025). Pyramidal neurons proportionately alter the identity and survival of specific cortical interneuron subtypes. bioRxiv.

46. Yadeta, K.A., and Thomma, B.P.H.J. (2013). The xylem as battleground for plant hosts and vascular wilt pathogens. Front Plant Sci 4, 97.

47. Yamada, T., Watanabe, S., Arai, M., Harada, K., and Kitamura, K. (2010). Cotyledonary node pre-wounding with a micro-brush increased frequency of Agrobacterium-mediated transformation in soybean. Plant Biotechnology 27, 4.

48. Yan, H., Lee, J., Song, Q., Li, Q., Schiefelbein, J., Zhao, B., and Li, S. (2022). Identification of new marker genes from plant single-cell RNA-seq data using interpretable machine learning methods. New Phytol 234, 1507–1520.

49. Yang, X., Zhao, W., Hua, C., Zheng, X., Jing, M., Li, D., Govers, F., Meijer, H.J., and Wang, Y. (2013). Chemotaxis and oospore formation in Phytophthora sojae are controlled by G-protein-coupled receptors with a phosphatidylinositol phosphate kinase domain. Mol Microbiol 88, 382–394.

50. Yu, G., Wang, L.G., Han, Y., and He, Q.Y. (2012). clusterProfiler: an R package for comparing biological themes among gene clusters. OMICS 16, 284–287.

51. Zhang, X., Luo, Z., Marand, A.P., Yan, H., Jang, H., Bang, S., Mendieta, J.P., Minow, M.A.A., and Schmitz, R.J. (2025). A spatially resolved multi-omic single-cell atlas of soybean development. Cell 188, 550–567 e519.

52. Zhang, Z., Gao, L., Ke, M., Gao, Z., Tu, T., Huang, L., Chen, J., Guan, Y., Huang, X., and Chen, X. (2022). GmPIN1-mediated auxin asymmetry regulates leaf petiole angle and plant architecture in soybean. J Integr Plant Biol 64, 1325–1338.

53. Zhang, Z., Luo, D., Zhong, X., Choi, J.H., Ma, Y., Wang, S., Mahrt, E., Guo, W., Stawiski, E.W., Modrusan, Z., Seshagiri, S., Kapur, P., Hon, G.C., Brugarolas, J., and Wang, T. (2019). SCINA: A Semi-Supervised Subtyping Algorithm of Single Cells and Bulk Samples. Genes (Basel) 10.

54. Zhou, F., Emonet, A., Denervaud Tendon, V., Marhavy, P., Wu, D., Lahaye, T., and Geldner, N. (2020). Co-incidence of Damage and Microbial Patterns Controls Localized Immune Responses in Roots. Cell 180, 440–453 e418.

55. Zhu, X., Fang, D., Li, D., Zhang, J., Jiang, H., Guo, L., He, Q., Zhang, T., Macho, A.P., Wang, E., Shen, Q.H., Wang, Y., Zhou, J.M., Ma, W., and Qiao, Y. (2023). Phytophthora sojae boosts host trehalose accumulation to acquire carbon and initiate infection. Nat Microbiol 8, 1561–1573.

