## Supplementary material for "Spatial Flavonoid Accumulation in Soybean Pericycle Restricts *Phytophthora sojae* Invasion": Figure S

**Figure S1. Early infection phenotypes of Wm and Wm82 at 4 h post-inoculation.**

**Figure S2. Cell Annotation assisted by Monocle and SCINA**

**Figure S3. Expression Patterns of Tissue-Specific Markers Used for SCINA-Assisted Cell Type Annotation.**

**Figure S4. Conserved infection-responsive genes exhibit amplified expression and broader cell-type specificity in resistant Wm82.**

**Figure S5. hdWGCNA reveals a Wm82 infection–associated gene co-expression module in pericycle.**

**Figure S6. Single-cell expression patterns of soybean flavonoid biosynthesis genes during *P. sojae* infection.**

**Figure S7. Generation and validation of *GmCHS7* and *GmCHI4A* knockout and overexpression soybean lines.**

**Figure S8. Effects of naringenin on *P. sojae* growth.**

**Figure S9. Dependence of *P. sojae* induced blue autofluorescence on flavonoids biosynthesis genes.**

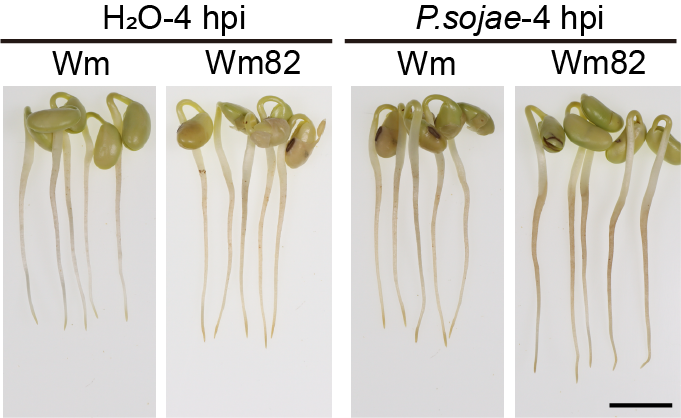

**Figure S1. Early infection phenotypes of Wm and Wm82 at 4 h post-inoculation.**

Wm and Wm82 seedlings were inoculated with *P. sojae* zoospores for 4 h (H₂O as control). Scale bar, 1 cm.

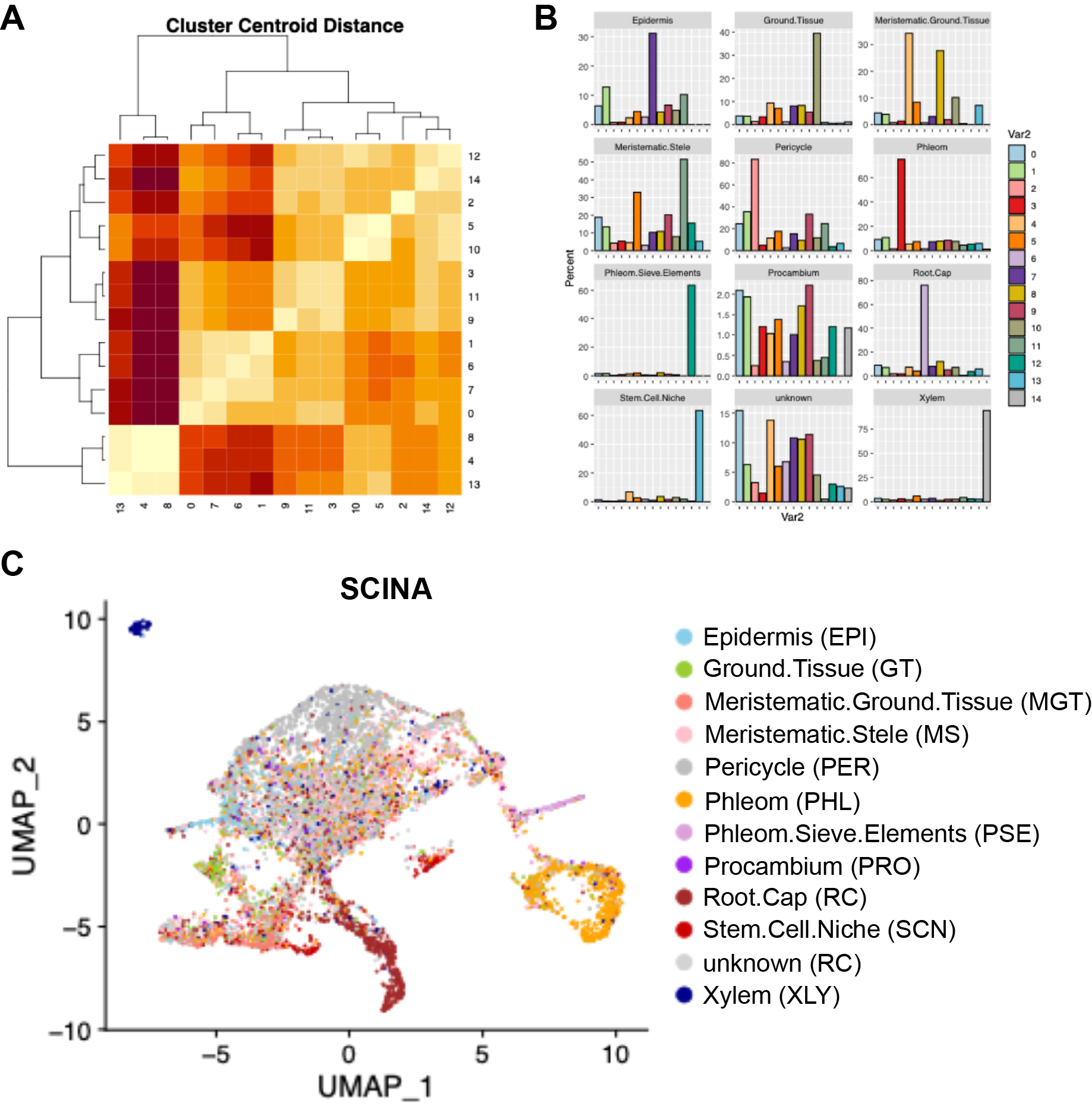

**Figure S2. Cell Annotation assisted by Monocle and SCINA**.

(A) Heatmap of Cluster Centroid Distance generated by Monocle2, showing the similarity between 15 clusters. The color gradient represents similarity, ranging from high (yellow) to low (dark red).

(B) Bar plot depicting cell identities assigned by SCINA for each individual cell type. The y-axis shows the percentage of cells within each cluster that are assigned a specific cell identity. (C) UMAP projection of cell identities assigned by SCINA, visualizing the spatial distribution of different cell types across the clusters.

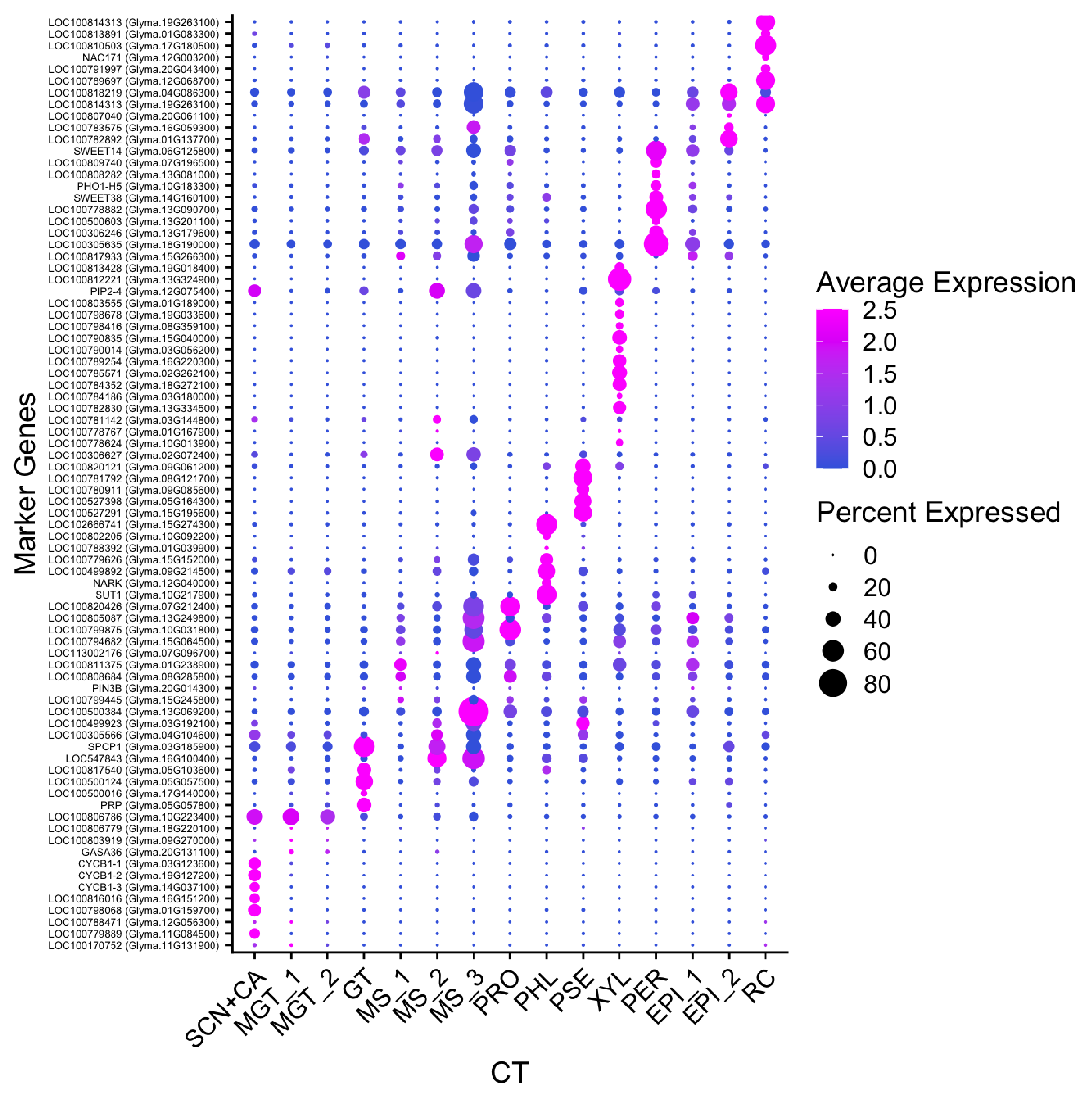

**Figure S3. Expression Patterns of Tissue-Specific Markers Used for SCINA-Assisted Cell Type Annotation.**

Bubble color indicates average gene expression intensity from low (blue) to high (magenta), while bubble diameter represents the percentage of cells expressing each gene within a given cluster.

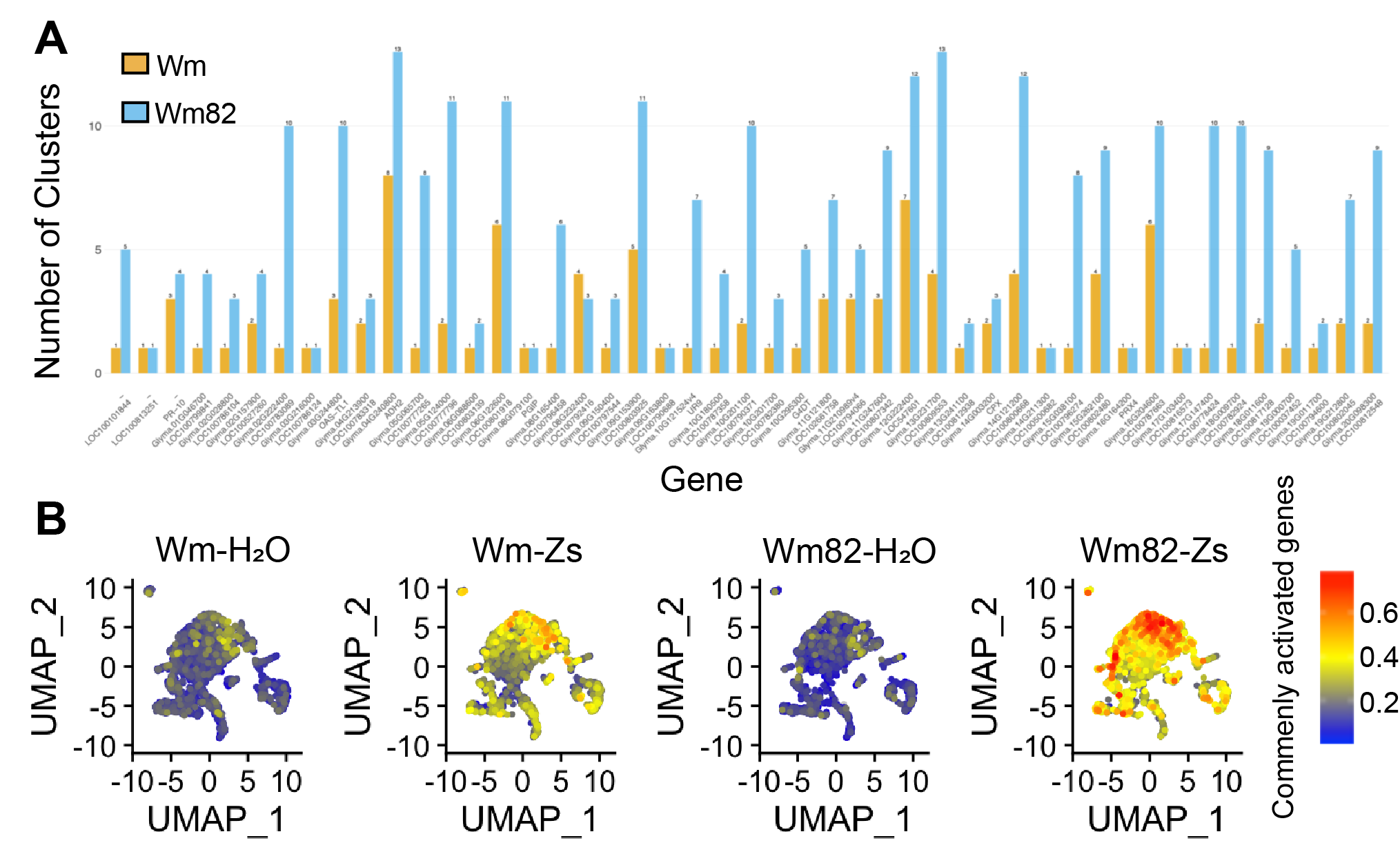

**Figure S4. Conserved infection-responsive genes exhibit amplified expression and broader cell-type specificity in resistant Wm82.**

**(A)** Barplot showing the number of cell clusters that a given gene is differentially expressed in Wm and Wm82.

**(B)** UCell score projection of commonly activated defense genes across samples from both genotypes. Color intensity represents expression level of the shared infection-responsive gene set, demonstrating significantly higher expression in Wm82 compared to Wm.

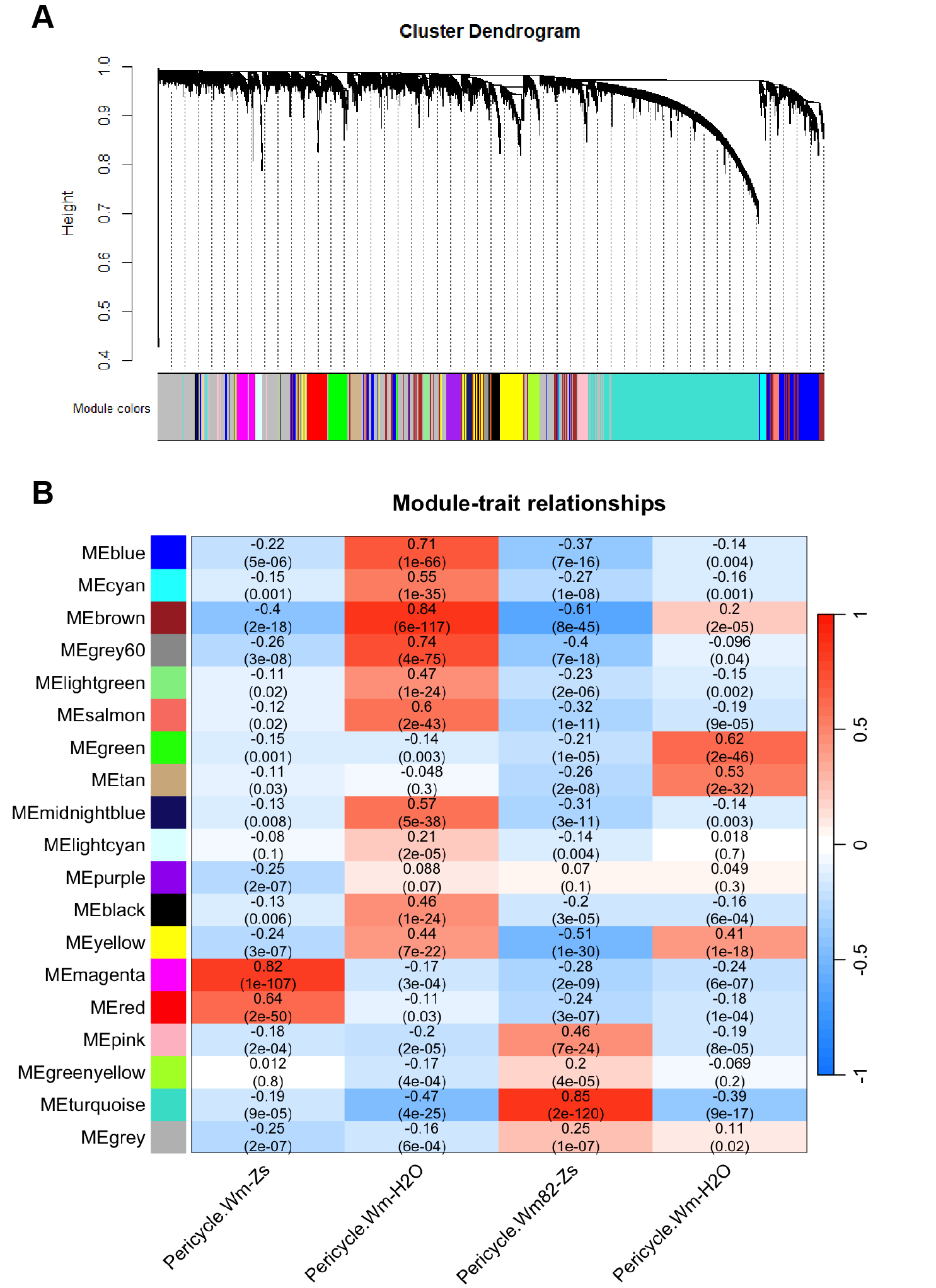

**Figure S5. hdWGCNA reveals a Wm82 infection–associated gene co-expression module in pericycle.**

**(A)** Hierarchical clustering dendrogram of gene co-expression modules identified by hdWGCNA in pericycle cells. Each branch represents a module of co-expressed genes, with colors indicating module identity. The largest module, designated the **Turquoise module**, contains the highest number of genes.

**(B)** Heat map of module–trait relationships showing correlations between each gene module and the four experimental conditions. The **Turquoise module** displays the strongest positive correlation with the **Wm82 post-infection (Wm82–Zs)** sample, suggesting that this module captures a core transcriptional program specifically activated during resistance.

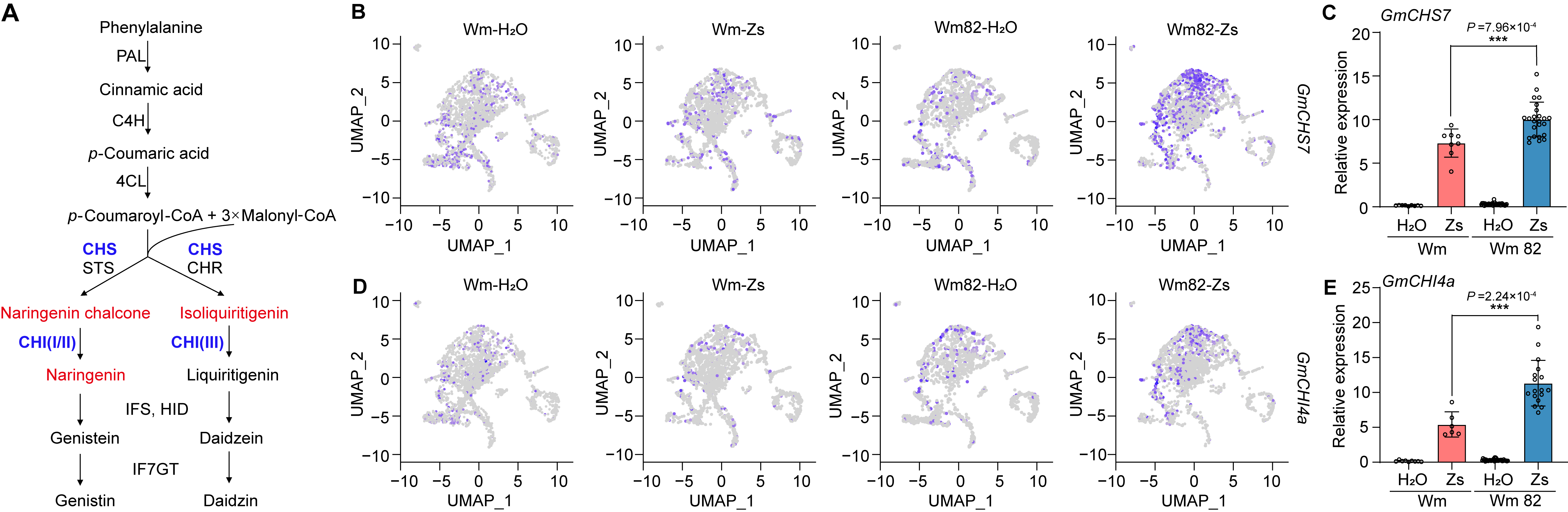

**Figure S6. Single-cell expression patterns of soybean flavonoid biosynthesis genes during *P. sojae* infection.**

(A) Schematic overview of the metabolic fluxes leading to isoflavonoid synthesis in soybean.

(B, D) UMAP projections showing the expression of *GmCHS7* and *GmCHI4A* in Wm82 under mock (H₂O) and *P. sojae* (Zs) conditions. Infection induces a clear shift in expression domains within the single-cell atlas.

(C, E) Relative expression levels of *GmCHS7* and *GmCHI4A* in Wm and Wm82 roots at 4 h post-inoculation with *P. sojae* zoospores. Both genes exhibit stronger induction specificly in the resistant genotype Wm82.

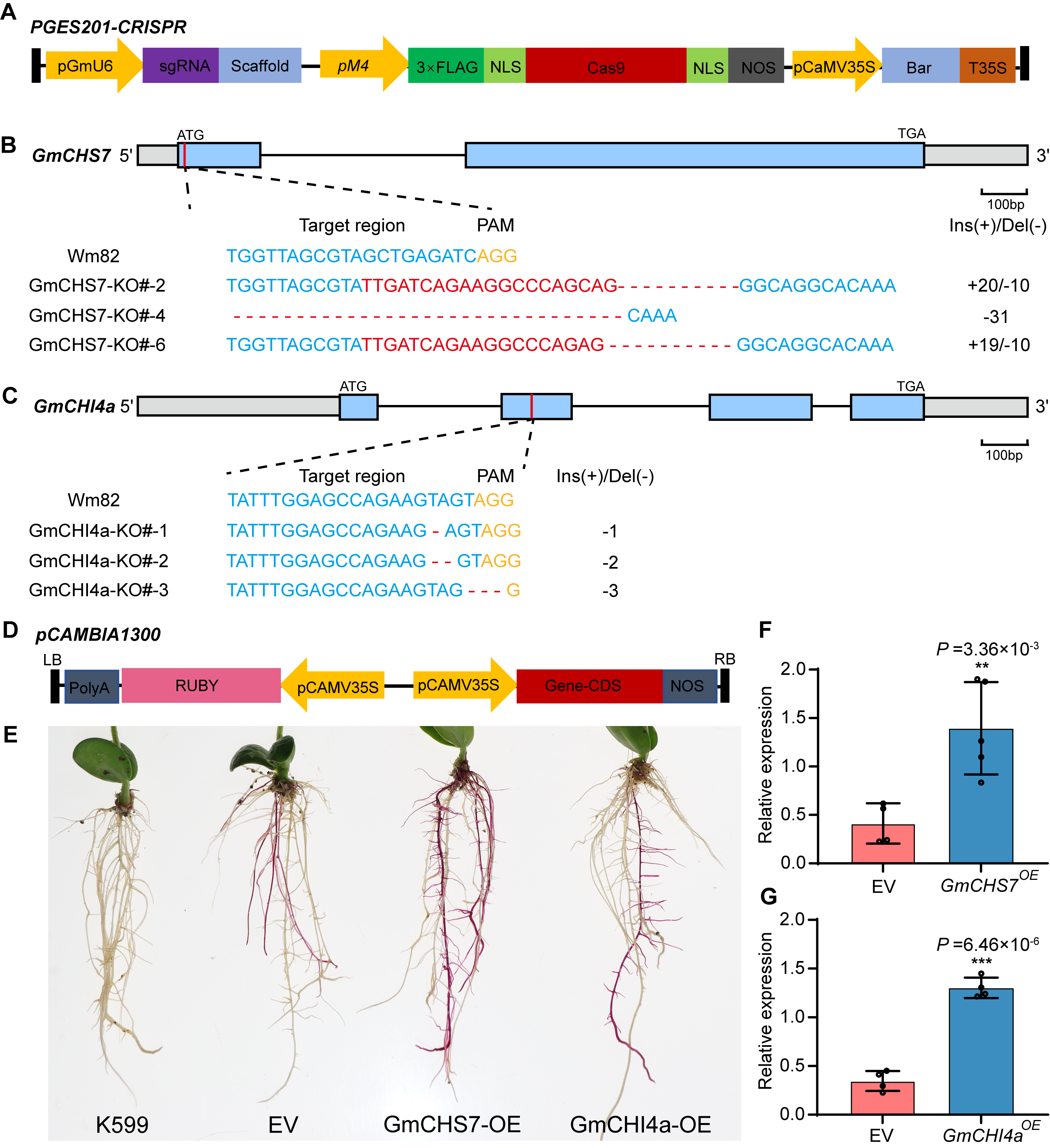

**Figure S7. Generation and validation of *GmCHS7* and *GmCHI4A* knockout and overexpression soybean lines.**

(A) Schematic showing the *PGES201* vector backbone used for CRISPR–Cas9–mediated gene knockout. Genes knockout were created by stable transformation.

(B, C) Gene-editing strategy and sgRNA target sites for generating *GmCHS7* and *GmCHI4A* knockout lines.

(D) Map of the *pCAMBIA1300*-based overexpression vector used to drive constitutive expression of target genes.

(E) Hairy-root transformation of soybean plants overexpressing *GmCHS7* and *GmCHI4A*.

(F, G) Relative expression levels of *GmCHS7* and *GmCHI4A* in transgenic roots, confirming successful knockout or overexpression in the generated lines.

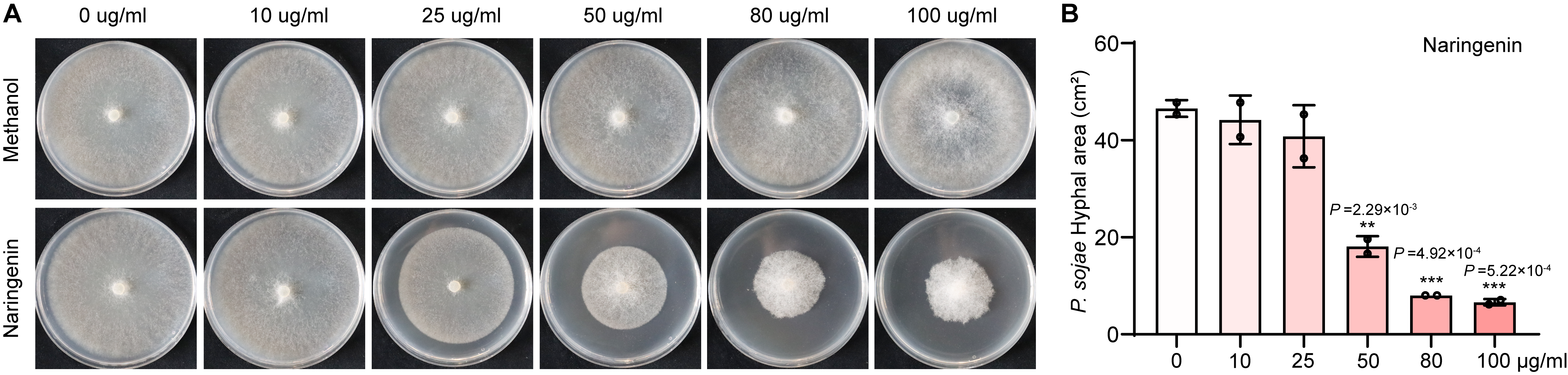

**Figure S8. Effects of naringenin on *P. sojae* growth.**

(A) Hyphal growth of *P. sojae* cultured for 5 days on plain 10% V8 or 10% V8 supplemented with methanol (solvent control) or naringenin dissolved in methanol at different concentrations.

(B) Quantification of hyphal growth on naringenin-containing 10% V8. The x-axis indicates naringenin concentration, and the y-axis shows the fungal growth area. Naringenin inhibits hyphal expansion in a dose-dependent manner.

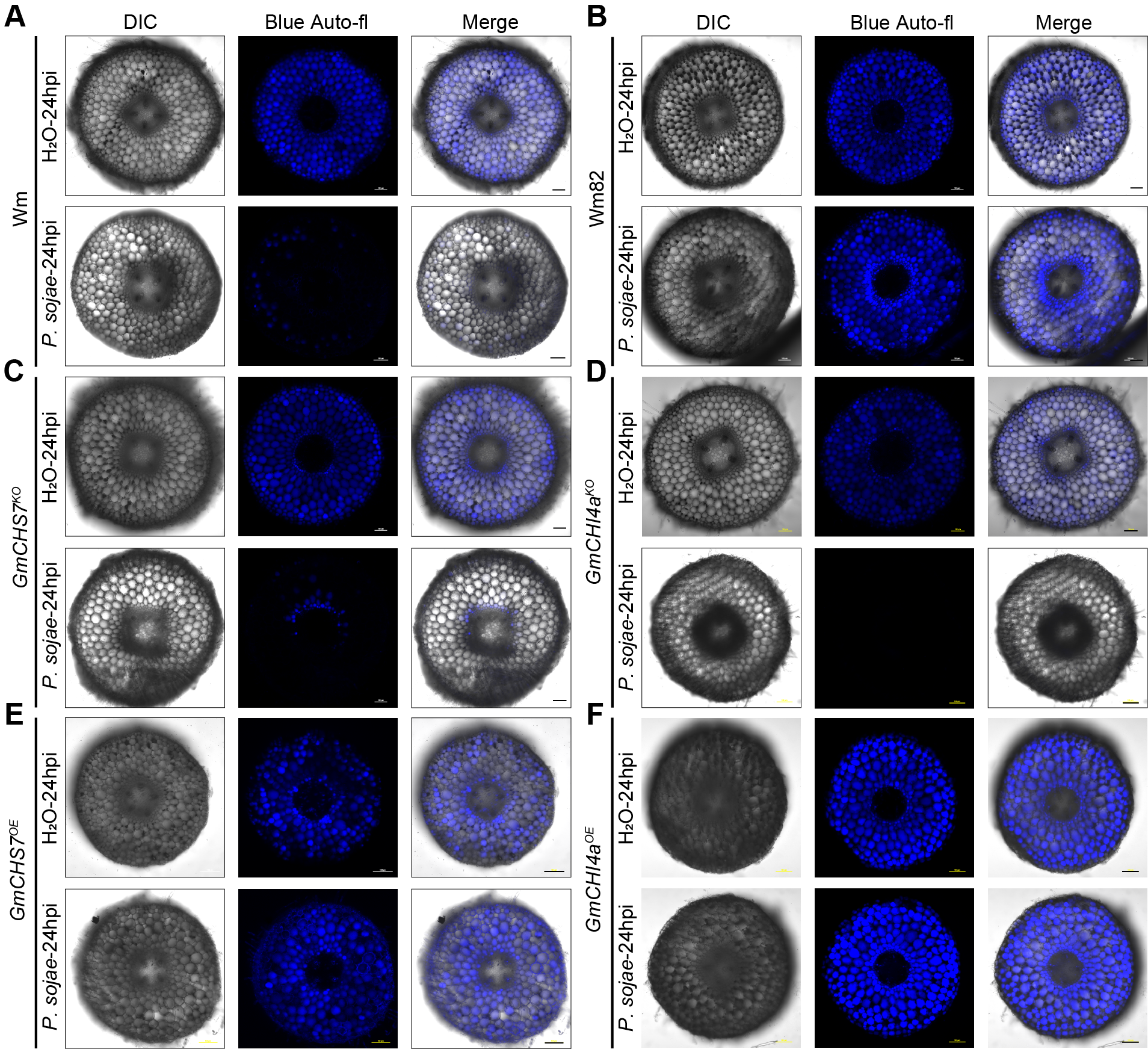

**Figure S9. Dependence of *P. sojae* induced blue autofluorescence on flavonoids biosynthesis genes.**

(A) Confocal microscopy images of root cross-sections from the susceptible cultivar Wm and at 24 hours post-inoculation (hpi) with *P. sojae*. In control conditions (H_2_O), blue autofluorescence is evenly distributed. Following infection, autofluorescence significantly diminishes in Wm.

(B) Confocal microscopy images of root cross-sections from the resistant cultivar Wm82 at 24 hours post-inoculation (hpi) with *P. sojae*. In control conditions, blue autofluorescence is evenly distributed. Following infection, autofluorescence exhibits a distinct autofluorescent band surrounding the stele, indicative of a localized immune response.

(C) Confocal microscopy images of root cross-sections from CRISPR knockouts of *GmCHS7* in the Wm82 genetic background, showing the absence of the autofluorescent band after *P. sojae* infection.

(D) Confocal microscopy images of root cross-sections from CRISPR knockouts of *GmCHI4A* in the Wm82 genetic background, showing the absence of the autofluorescent band after *P. sojae* infection.

(E) Confocal microscopy images of root cross-sections from *GmCHS7* overexpression lines in the Wm background, showing ubiquitous blue autofluorescence under both control and infection conditions.

(F) Confocal microscopy images of root cross-sections from *GmCHI4A* overexpression lines in the Wm background, showing ubiquitous blue autofluorescence under both control and infection conditions. Scale bar, 100 μm
